# Efficacy of plasmid-encoded CRISPR-Cas antimicrobial is affected by competitive factors found in wild *Enterococcus faecalis* isolates

**DOI:** 10.1101/2022.03.08.483478

**Authors:** Dennise Palacios Araya, Moutusee Islam, Shah O. Moni, Christine A. Ramjee, Tuong-Vi Cindy Ngo, Kelli L. Palmer

**Affiliations:** Department of Biological Sciences, University of Texas at Dallas, Richardson, TX, USA, 75080

**Author notes:** Correspondence: Kelli L. Palmer.

**Keywords:** *Enterococcus faecalis*, CRISPR-Cas, antibiotic resistance, CRISPR-Cas antimicrobial, model strain, wild isolate, cytolysin, Bac-21

## Abstract

*Enterococcus faecalis* is a leading cause of hospital-acquired infections. These infections are becoming more difficult to treat due to the increasing emergence of *E. faecalis* strains resistant to last resort antibiotics. Over the past decade, multiple groups have engineered the naturally occurring bacterial defense system CRISPR-Cas as a sequence-specific antimicrobial to combat antibiotic-resistant bacteria. We have previously established that the type II CRISPR-Cas system of *E. faecalis* can be reprogrammed as a CRISPR-Cas antimicrobial and delivered to antibiotic-resistant recipients on a conjugative pheromone-responsive plasmid. Using a co-culture system, we showed sequence-specific depletion of antibiotic resistance from *E. faecalis* model strains, both *in vitro* and *in vivo*. Although this and other studies have demonstrated the potential use for CRISPR-Cas as an antimicrobial, most have deployed the system against model bacterial strains. Thus, there is limited knowledge on how effective these potential therapies are against recently isolated and uncharacterized strains with limited laboratory passage, which we refer to here as wild strains. Here, we compare the efficacy of our previously established CRISPR-Cas antimicrobials against both *E. faecalis* model strains and wild *E. faecalis* fecal isolates. We demonstrate that these wild isolates can antagonize the CRISPR-Cas antimicrobial donor strain via competitive factors like cytolysin. Furthermore, we show that the wild isolates can effectively prevent delivery of the CRISPR-Cas antimicrobial plasmids, consequently avoiding CRISPR-Cas targeting. Our results emphasize the requisite to study CRISPR-Cas antimicrobials against wild strains to understand limitations and develop delivery systems that can endure competitive interspecies interactions in the gut microenvironment and effectively deliver CRISPR-Cas antimicrobials to their intended targets.

**IMPORTANCE:** *Enterococcus faecalis* is a major nosocomial pathogen. Traditional antibiotics continue to lose potency against these opportunistic pathogens as they become increasingly resistant to more antibiotics. We previously showed that our CRISPR-Cas antimicrobials can deplete drug resistance in or kill *E. faecalis* model strains. Here, we examined the efficacy of CRISPR-Cas antimicrobials against a recent collection of *E. faecalis* fecal isolates. We found that CRISPR-Cas delivery and efficacy is affected by competitive factors produced by the wild isolates. Our study emphasizes the need to study CRISPR-Cas antimicrobials in the context of wild bacterial isolates, which are the intended target for this potential therapy, in order to understand limitations and develop CRISPR-enhanced antimicrobials with effective clinical applications.

## INTRODUCTION

The rapid emergence and spread of drug-resistant pathogens are leading global health concerns in the 21^st^ century (1). The overuse of antibiotics in medical and agricultural settings and the acquisition and dissemination of mobile genetic elements (MGEs) are significant contributors to the emergence of multidrug-resistant (MDR) pathogens (2). *Enterococcus faecalis* is a common commensal in the gut of healthy humans (3), however, under certain circumstances, i.e., in immunocompromised people or under a dysbiotic environment, they can act as opportunistic pathogens (4, 5). *E. faecalis* has emerged as a major cause of hospital-acquired infections (3). Importantly, these infections are becoming harder to treat as more *E. faecalis* strains become resistant to antibiotics such as vancomycin, β-lactams, and aminoglycosides (3, 6). The limited discovery and development of new antimicrobial drugs points to the need for alternative therapies to combat drug-resistant *E. faecalis*.

The past decade has been fundamental in the exploration of novel therapeutic approaches to control bacterial infections. Pioneering studies have demonstrated that engineered CRISPR-Cas systems can be used as sequence-specific antimicrobials to target the bacterial chromosome, thereby killing or inhibiting pathogens, or to eliminate plasmids harboring drug-resistance genes (7, 8). CRISPR-Cas are natural defense mechanisms that work as adaptive immune systems against MGEs (9). For Type II systems, CRISPR-Cas targeting activity is achieved by the endonuclease Cas9, which forms a complex with a guide-RNA (gRNA). Upon encountering a MGE with complementary sequence to the gRNA, the Cas9/gRNA complex generates a double-stranded break in the MGE, causing sequence-specific elimination of the invading element (9).

A critical aspect of adapting CRISPR-Cas systems as antimicrobials is the development of robust delivery vectors. One delivery approach is to use modified phages such as phagemids (7, 10–12). Although phage-mediated delivery has been found to be very robust, phage rely on cell receptors which are a common mechanism for bacterial resistance (13). Conjugative plasmid delivery of CRISPR-Cas antimicrobials has also been proposed for their flexible host ranges and cellular receptor independence (13–15).

In an earlier study, the natively encoded *E. faecalis* Cas9 was co-expressed with a self-targeting CRISPR array and deployed using mobilizable cloning plasmids (14, 16). It was determined that the Type II CRISPR-Cas system of *E. faecalis* could be utilized as an effective, programmable tool to alter *E. faecalis* populations (16). To explore this concept further, we previously designed a CRISPR-Cas antimicrobial plasmid that uses the pheromone-responsive plasmid (PRP) pPD1 as the delivery backbone (14). This plasmid was chosen because of its high conjugation frequency, native lack of antibiotic resistance genes, and its self-selection and fitness-enhancing features, which are conferred by the bacteriocin Bac-21 encoded within pPD1 (17, 18). The engineered pPD1 plasmid, termed pKH88, encodes a constitutively expressed CRISPR array/Cas9 module programmed to target either the *tetM* gene (encoding tetracycline resistance) or the *ermB* gene (encoding erythromycin resistance). Using both *in vitro* and *in vivo* mouse colonization models, we demonstrated the ability of these CRISPR-Cas antimicrobials to deplete antibiotic resistance from *E. faecalis* oral and clinical isolates commonly utilized for laboratory studies (14).

These results led us to investigate whether our engineered CRISPR-Cas antimicrobials are effective against a recent collection of human fecal surveillance *E. faecalis* isolates (19). Previous to this study, this *E. faecalis* collection had not been extensively characterized nor fully sequenced, hence we considered these isolates uncharacterized. Moreover, because this is a recent collection with limited laboratory handling and manipulation, we also considered them “undomesticated” or ‘wild’ isolates. We explored this question because despite the extensive research in this area, to our knowledge, only one study has examined the efficacy of CRISPR-Cas antimicrobials using uncharacterized, wild strains (20). Instead, many studies have focused primarily on investigating CRISPR-Cas antimicrobial targeting in well-characterized model strains (11–14, 21).

Here, we compare the efficacy of our CRISPR-Cas antimicrobials against both *E. faecalis* model strains and *E. faecalis* uncharacterized, wild isolates. We show that the *E*. *faecalis* wild isolates are adept at competing against the CRISPR-Cas antimicrobial donor strain. This is in part due to production of competitive factors such as the enterococcal cytolysin, a two-component peptide toxin that is commonly found in highly virulent *E. faecalis* strains and that has hemolytic activity and bactericidal activity against many Gram-positive bacteria (22–24). We also show that the wild isolates can successfully prevent mobilization of the CRISPR-Cas antimicrobial plasmids, particularly the ones carrying a cognate guide-RNA, by an unknown mechanism. Importantly, we demonstrate that inefficient delivery of the CRISPR-Cas plasmids prevents sequence-specific depletion of antibiotic resistance. Notably, none of these aforementioned phenotypes were observed among the model strains, which may be due to laboratory domestication causing loss of competitive fitness and defense mechanisms as previously shown for other bacteria (25–29). The challenges that we encountered with our *E. faecalis* wild isolates emphasizes the need to further explore CRISPR-Cas antimicrobials in the context of wild strains, as other delivery vectors may encounter similar roadblocks. Understanding potential limitations will ensure that we engineer delivery vectors (i.e., plasmid donor strains if using conjugative plasmids) to withstand the harsh gut microenvironment and successfully deliver the CRISPR-Cas antimicrobial.

## RESULTS

### Chromosomal targeting of *vanB* in the MDR *E. faecalis* strain V583 results in depletion of both vancomycin resistance and viable V583 cells from the population

We have previously reported that the CRISPR-Cas antimicrobial plasmids pKH88[sp-*tetM*] and pKH88[sp-*ermB*] have the ability to selectively remove tetracycline and erythromycin resistance, respectively, from the *E. faecalis* model strain OG1SSp. In that study, OG1SSp harbored either pCF10, carrying tetracycline resistance gene *tetM*, or pAM771, carrying erythromycin resistance gene *ermB.* It was further established that delivery of pKH88[sp-*ermB*] was able to kill the *E. faecalis* model strain V583 (14), a vancomycin-resistant hospital-adapted isolate (30), by targeting *ermB* on plasmid pTEF1.

Plasmids pKH88[sp-*tetM*] and pKH88[sp-*ermB*] derive from the pheromone-responsive plasmid pPD1 (14). These derivatives constitutively express the *cas9* gene and an engineered CRISPR guide RNA. To achieve transconjugant detection, a chloramphenicol resistance marker (*cat*) was inserted into the plasmid. These derivatives retain pheromone response and production of bacteriocin 21 (Bac-21), which is natively encoded on pPD1 (Figure 1A) (14, 31).

**Figure 1.**
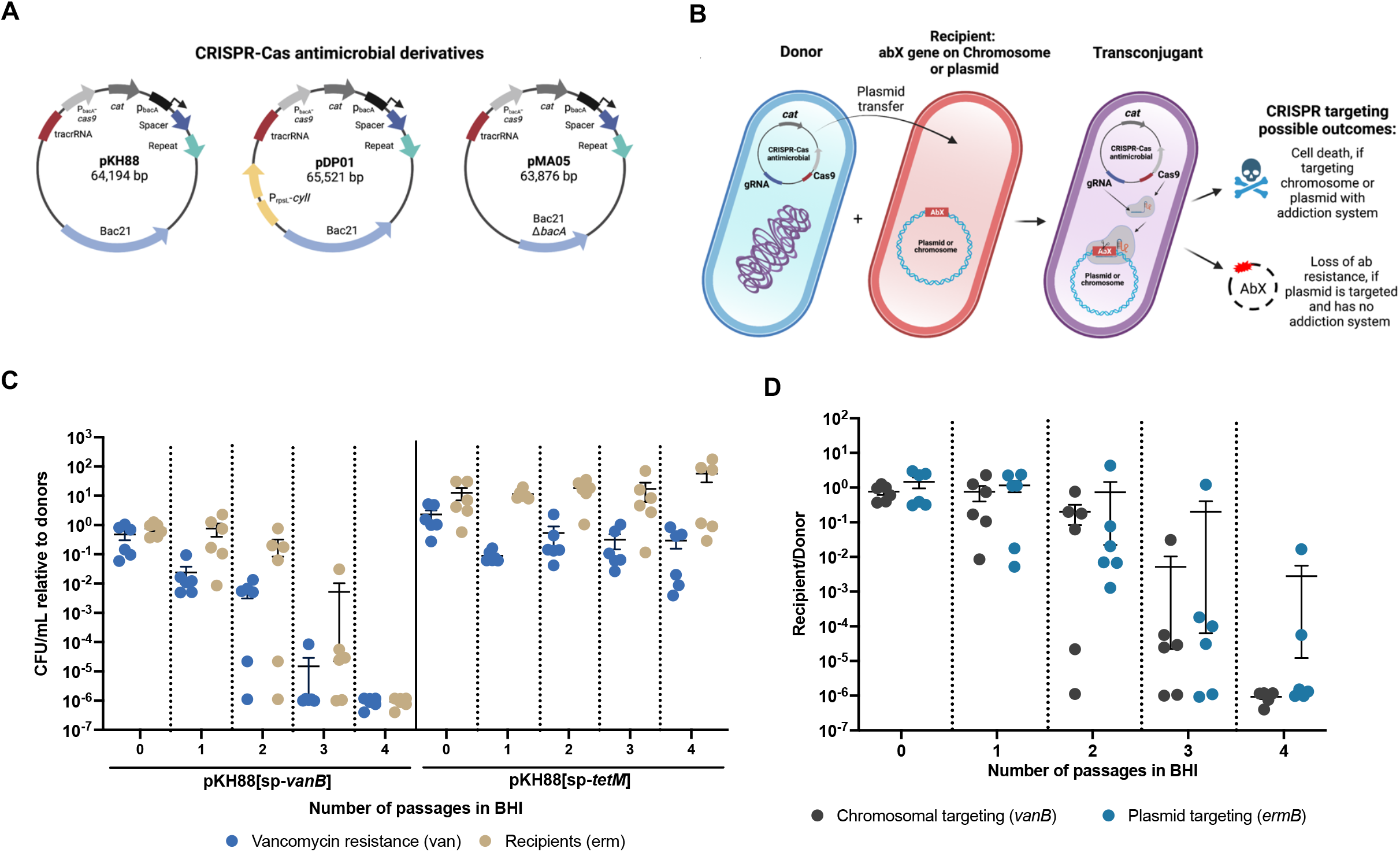
Conjugative delivery of engineered CRISPR-Cas antimicrobial plasmids to eliminate antibiotic resistance from *E. faecalis* using an *in vitro* co-culture system. (A) Schematic of the pKH88 CRISPR-Cas antimicrobial plasmid derivatives, which include adding the cytolysin immunity gene *cylI* (pDP01) or deleting the bacteriocin 21 precursor gene *bacA* from the pPD1 backbone (pMA05). Plasmids encode Cas9 and single-guide RNA sequences to target antibiotic resistance genes *tetM*, *ermB* or *vanB.* (B) Experimental design of the CRISPR-antimicrobial assays. (C) Vancomycin resistant cells and V583 recipients (erythromycin-resistant cells) are depleted from the population when pKH88[sp-*vanB*] donors are present, which target the *vanB* gene on the V583 chromosome. No comparable depletion is observed when donors with the nontargeting plasmid pKH88[sp-*tetM*] are present. (D) No statistically significant difference was observed when targeting *vanB* on the V583 chromosome versus targeting *ermB* on V583 plasmid pTEF1 at any timepoint. Error bars represent the mean plus the standard error of the mean (SEM). Mann-Whitney test; P-value: P0: 0.937, P1: 0.818, P2: 0.937, P3: 0.588, P4: 0.060.

To gain further insight into the response of V583 when targeted at different locations, we engineered a third CRISPR-Cas antimicrobial, termed pKH88[sp-*vanB*], which encodes a guide-RNA targeting the vancomycin resistance gene *vanB* present on the chromosome of V583 (24). V583 does not encode *tetM* (24), therefore pKH88[sp-*tetM*] was used as a nontargeting control. The *E. faecalis* strain CK135RF, encoding resistance to rifampicin and fusidic acid, was used as the plasmid donor (Figure 1B). We tested whether targeting the chromosome of V583 would elicit a different response (i.e., faster removal of resistant bacteria) than targeting its plasmid pTEF1. As expected, chromosomal targeting resulted in decreased *E. faecalis* V583 cells as well as depletion of total vancomycin resistance from the population (Figure 1C). Remarkably, reduction of viable V583 cells happened at a similar rate compared to targeting plasmid pTEF1 (Figure 1D). These results support the notion that while in most cases CRISPR cleavage of plasmids cause their loss (7, 10), in some instances, the loss of the plasmid results in cell death (14).

### Screening a collection of *E. faecalis* fecal isolates for CRISPR-Cas antimicrobial targeting

In a practical setting, CRISPR-Cas antimicrobials would be utilized to either eliminate drug-resistance genes from uncharacterized *E. faecalis* strains or to remove resistant *E. faecalis* from a community altogether. Therefore, in this study we evaluated whether removal of the resistant bacteria and/or sequence-specific depletion of antibiotic resistance could be achieved in a recent collection of *E. faecalis* fecal surveillance isolates using our developed CRISPR-Cas antimicrobials (14, 19).

We began by screening the fecal isolates by PCR to confirm the presence of genes *tetM* and *ermB* (Table 1), targeted by pKH88[sp-*tetM*] and pKH88[sp-*ermB*], respectively. This initial genotyping was later supplemented by whole genome sequencing to confirm results (Table 1 and described further below). The protospacer (i.e., sequence being targeted by CRISPR-Cas) and required NGG protospacer adjacent motif (PAM) sequences of *ermB* and *tetM* were verified to confirm 100% match with the targeting sequences selected for the CRISPR-Cas antimicrobials (Table S1). Although isolate 2-1 encodes *tetM* and presents a resistant phenotype, an NGG PAM sequence was not present next to the protospacer in *tetM*, hence testing pKH88[sp-*tetM*] on isolate 2-1 was excluded (Table S1). We had previously determined that none of the fecal isolates harbor the *vanB* gene (19), thus pKH88[sp-*vanB*] was used as a nontargeting control.

**Table 1.** Qualitative analysis of Enterococcus faecalis fecal isolates.

| Isolate | *cylA* |  | Hemolysis assay | *tetM* |  | *ermB* |  | cPD1 clumping assay |
| --- | --- | --- | --- | --- | --- | --- | --- | --- |
|  | PCR | Sequence analysis |  | PCR | Sequence analysis | PCR | Sequence analysis |  |
| 2-1 | + | + | + | - | + | + | + | + |
| 43-2 | + | + | + | + | + | + | + | - |
| 59 | + | + | ♀ | + | + | + | + | - |
| 94-1 | + | + | + | - | + | + | + | + |
| 101-1 | - | - | - | + | + | + | + | - |
| 119-1 | + | + | + | - | + | + | + | - |
| 119-2 | + | + | + | - | + | + | + | - |
| 133-1 | - | - | - | + | + | + | + | - |
| 142-1 | - | - | - | + | + | + | + | - |
| 143-1 | + | + | + | - | - | + | + | - |
| 152 | + | + | + | - | + | + | + | - |

*cylA*, gene encoding the activator component in the cytolysin operon; *tetM*, gene encoding tetracycline resistance; *ermB*, gene encoding erythromycin resistance; cPD1, sex pheromone that pPD1 responds to.
♀ Inconclusive results.
\* Isolate 2-1 clumps in the absence of pheromone.

We also generated an antibiotic resistance profile to determine which isolates met the criteria for conjugation assays. Specifically, the selected isolates must have tetracycline and/or erythromycin resistance phenotypes to be targeted, no chloramphenicol-resistant phenotype, since this antibiotic was used as a selective marker for the CRISPR-antimicrobial plasmids, and no rifampicin- and fusidic acid-resistant phenotype, since the CRISPR-antimicrobial plasmid donor had those markers (Table 2). Furthermore, since we needed a universal marker to select for the fecal isolates (recipients), we also screened for resistance to streptomycin. Three isolates were found to be resistant or partially resistant to streptomycin (Table 2). We selected for spontaneous streptomycin-resistant mutants in each of the remaining isolates. Additionally, because these CRISPR-Cas antimicrobials derive from pPD1 (14), we also performed pheromone response assays with synthetic cPD1 pheromone to assess whether these fecal isolates already carried pPD1 (i.e., exhibited cell clumping in response to cPD1), and thus potentially would not uptake the CRISPR-Cas antimicrobials (Table 1).

**Table 2.** *Enterococcus faecalis* fecal isolate growth on antibiotic-containing agars.

| Fecal isolate | Antibiotic |  |  |  |  |  |
| --- | --- | --- | --- | --- | --- | --- |
|  | Tet | Erm | Van | Cm | RF | Stm |
| 2-1 | G | G | - | - | - | - |
| 43-2 | G | G | - | - | - | G* |
| 59 | G | G | G | - | - | - |
| 94-1 | G | G | - | - | - | - |
| 101-1 | G | G | - | - | - | G |
| 119-1 | G | G | - | - | - | - |
| 119-2 | G | G | - | - | - | - |
| 133-1 | G | G | - | - | - | G |
| 142-1 | G | G | - | - | - | - |
| 143-1 | - | G | - | - | - | - |
| 152 | G | G | - | - | - | - |

Tet, tetracycline; Erm, erythromycin; Van: vancomycin; Cm, chloramphenicol; RF, rifampicin and fusidic acid; Stm, streptomycin; G: Growth; Dash (-): No growth; G\*: Growth but colonies smaller than wild-type colonies after 24 h of incubation. See materials and methods for antibiotic concentrations used.

Based on the screenings, we chose 11 fecal isolates from our collection, all of which harbor the *ermB* gene, while 10 also carry the *tetM* gene (Table 1). All isolates grew on erythromycin and/or tetracycline supplemented plates, confirming that the resistance genes were expressed (Table 2). Furthermore, 9 of the isolates did not clump in the presence of synthetic pheromone while 2 had clumping phenotypes (Table 1). Additional assessment of those two isolates through genomic sequence analysis confirmed that they do not harbor plasmid pPD1, thus were appropriate for further testing (Table S1). Table 1 shows the list of fecal isolates that were determined to be suitable to examine targeting by our CRISPR-Cas antimicrobials.

### Delivery of CRISPR-Cas antimicrobials is impaired by low donor survival and low conjugation frequencies

Next, we performed conjugation assays using *E. faecalis* CK135RF as the CRISPR-Cas antimicrobial donor strain, and each individual *E. faecalis* fecal isolate as recipients (Figure 1B). As controls, we also performed assays using the *E. faecalis* model strain OG1SSp possessing either pCF10 (carrying the *tetM* gene) or pAM714 (carrying the *ermB* gene) as recipients (32). Donors and recipients were mixed in a 1:10 ratio, plated on BHI medium without antibiotics and incubated overnight (passage 0, P0). The biofilm was harvested, diluted, and used as inoculum in liquid medium. The bacterial community was allowed to grow for a second overnight in this medium (passage 1, P1). Following each incubation, the CFU/mL of specific populations (e.g., donor, transconjugant, et cetera) was determined.

After conjugation, sequence-specific depletion of antibiotic resistance was established as follows: We first calculated the average CFU/mL recovered from erythromycin and tetracycline-supplemented plates after treatment. This was determined for both targeted and nontargeted populations. Then, the average CFU/mL of the targeted population was divided by the average CFU/mL of the nontargeted population. Calculations were computed for both P0 and P1. If the value was ≥1, then no sequence-specific depletion was deemed to have occurred. If the value was ≤0.1, then at least 10-fold sequence-specific depletion in resistance was deemed to have occurred. Based on these parameters, only the controls and fecal isolates 59 and 101-1 showed sequence-specific decreases in tetracycline and erythromycin resistance at passage 1. Isolate 59 revealed only partial decrease for both antibiotics (∼10 to 30-fold change) while isolate 101-1 showed between ∼230 to 480-fold change, although neither was statistically significant (Figure 2). We also observed a partial decrease of both erythromycin and tetracycline resistance in isolate 133-1 (Figure 2) and of erythromycin resistance in isolate 142-1 (Figure 2A), although not in a sequence-specific manner since the reduction was also observed when challenging the cells with the nontargeting plasmid. In addition, fecal isolates 59 and 101-1 showed a reduction in recipient cell numbers regardless of the location of the targeted gene (plasmid or chromosome) (Figure S1 and Table S1). These results are consistent with our previous observation that cleavage of *E. faecalis* V583 pTEF1 causes cell death (14).

**Figure 2.**
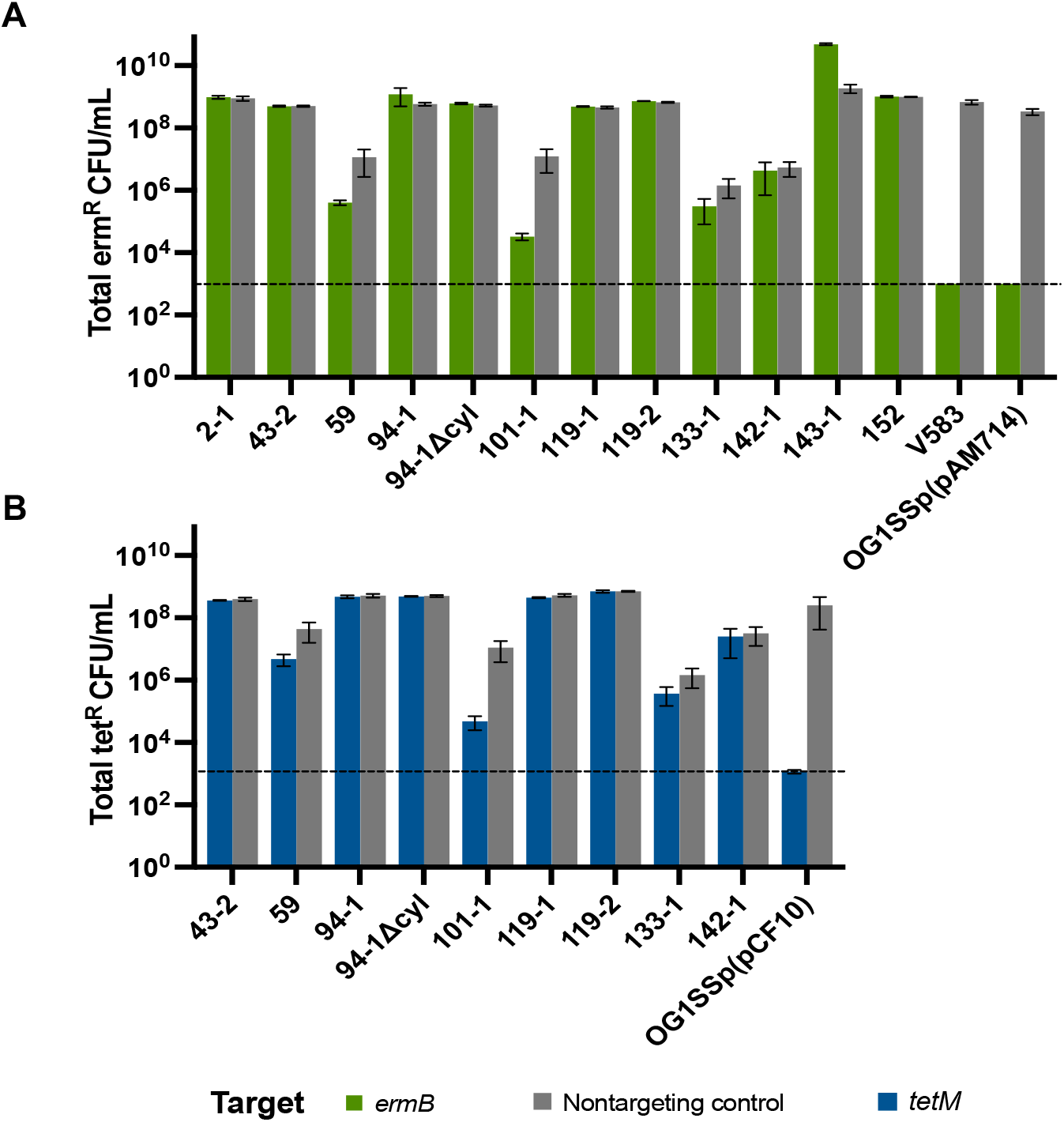
Total antibiotic resistance remaining in the population at passage 1 when using pKH88 derivatives as CRISPR-Cas antimicrobials. Sequence-specific depletion of erythromycin or tetracycline resistance was observed when targeting the (A) *ermB* or (B) *tetM* gene in the model strains V583 (encodes *ermB*), OG1SSp:pAM714 (encodes *ermB*) and OG1SSp:pCF10 (encodes *tetM*), respectively. Targeting of the fecal isolates did not result in substantial resistance reduction. A small sequence-specific depletion of both erythromycin and tetracycline resistance was observed for isolates 59 and 101-1, although not statistically significant (unpaired t-test; P-value >0.2). Error bars represent the geometric mean and SEM of data from three independent biological experiments.

Unexpectedly, the donor population was near or below the level of detection (1×10^3^) in experiments with seven of the fecal isolates (Figure 3, blue bars and Figure S2), indicating that the donors competed poorly against those isolates. Moreover, conjugation of the nontargeting plasmid was observed in 10 of the 11 fecal isolates (as measured by transconjugant CFU/mL), while conjugation of the targeting plasmids was less frequent (only detected in 5 isolates) and transconjugants were at lower densities, near or below the level of quantification (2.5×10^4^ CFU/mL) (Figure 4A).

**Figure 3.**
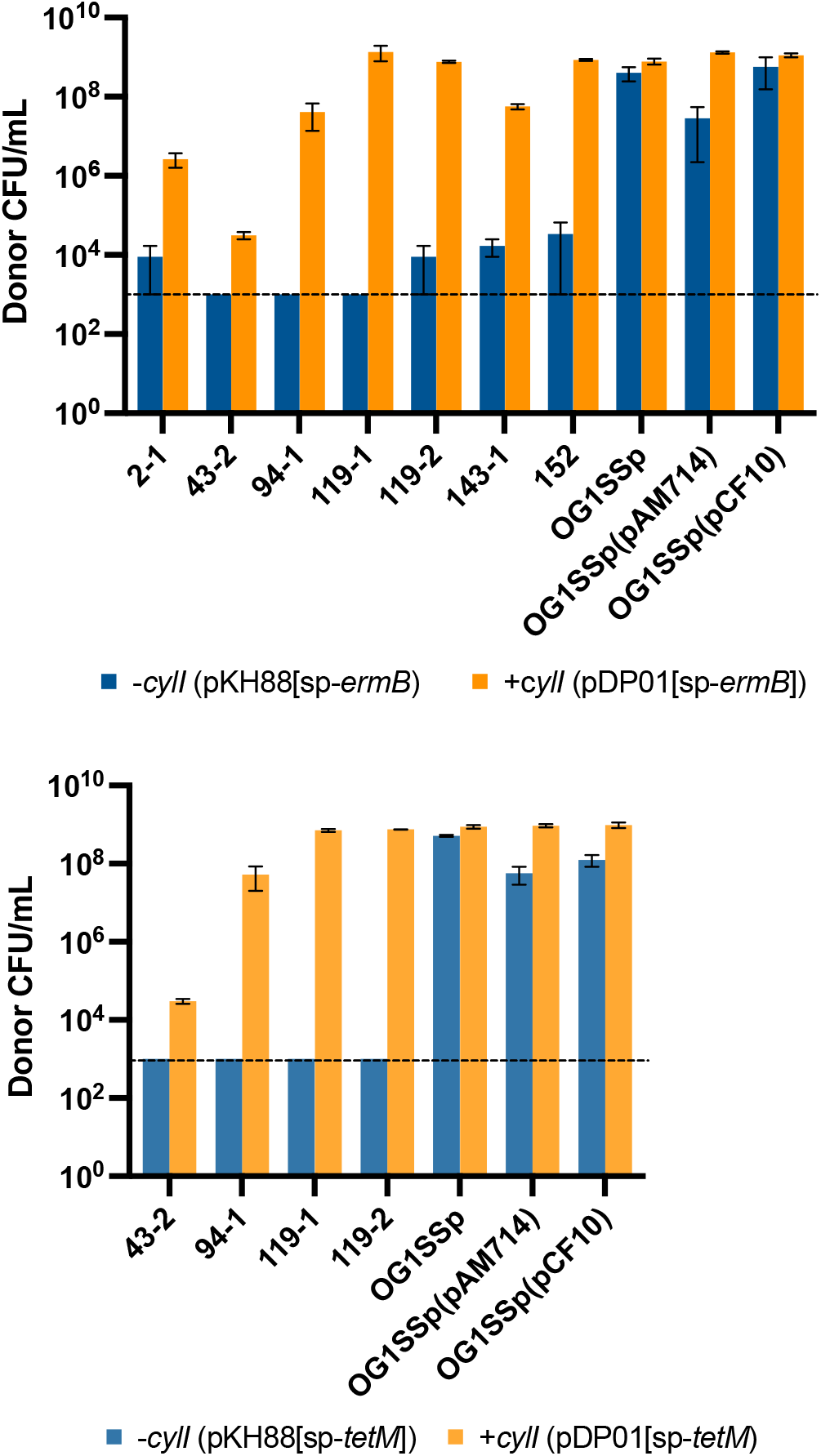
Survival of the CRISPR-antimicrobial donor harboring pKH88 or pDP01 derivatives at passage 1. Donors carrying the pKH88 plasmids competed poorly against cytolysin-positive fecal isolates (blue bars), while donors carrying the pDP01 plasmids (encoding cytolysin immunity) had higher survival rates (orange bars). Donors competing against model strains survived well regardless of the CRISPR-Cas antimicrobial plasmid used. Error bars represent the geometric mean and SEM of data from three independent biological experiments.

**Figure 4.**
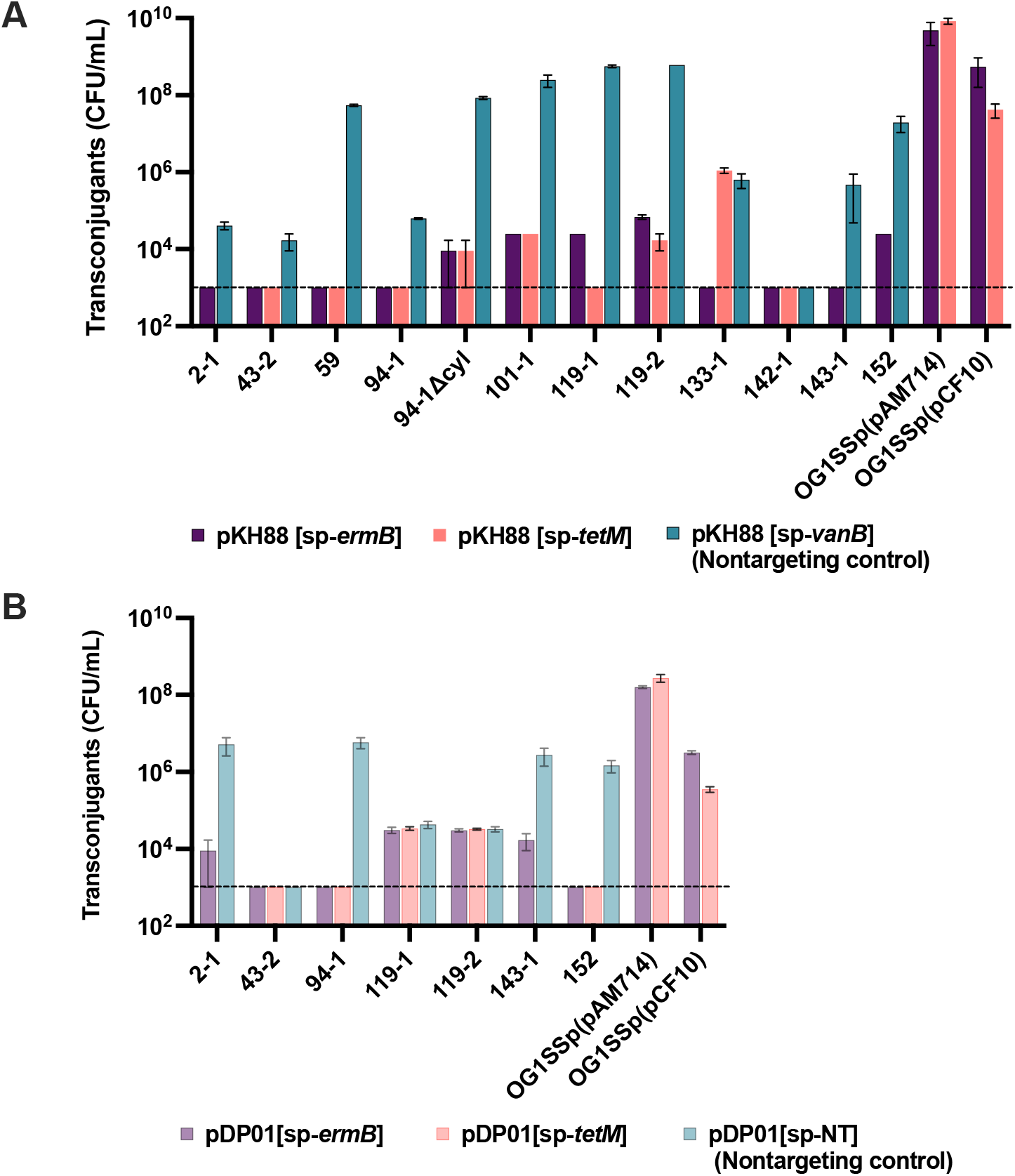
Conjugation of CRISPR-Cas antimicrobial plasmids into different *E. faecalis* strains at passage 0. (A) Low transconjugant numbers were obtained when treating the fecal isolates with pKH88 targeting plasmids while treatment with the pKH88 nontargeting plasmid resulted in most isolates acquiring the plasmid. While the model strains OG1SSp(pAM714) and OG1SSp(pCF10) were able to acquire both the targeting and nontargeting plasmids at similar rates. (B) Similarly, low transconjugant numbers were observed when treating the fecal isolates with pDP01 (harboring cytolysin immunity). The model strains acquired the pDP01 plasmids at similar or higher rates than the fecal isolates. Error bars represent the geometric mean and SEM of data from three independent biological experiments.

In contrast to these observations, identical conjugation assays using the well-characterized *E. faecalis* model strain OG1SSp carrying pCF10 or pAM714 as the recipient resulted in high donor survival (Figure 3 and S2), a high number of transconjugants of both targeting and nontargeting plasmids (Figure 4A), and reduction in erythromycin and tetracycline resistance below the limit of detection when receiving a pKH88 derivative encoding a cognate RNA-guide (Figure 2).

Based on our results, we hypothesized that there is a sequence-specific effect on the entry of the CRISPR-antimicrobial plasmids into the fecal isolates. To test this hypothesis, we circumvented conjugation by directly transforming plasmids into electrocompetent cells. For these experiments, we utilized vectors that were used to integrate the CRISPR-Cas machinery into plasmid pPD1 and generate the pKH88 derivatives (14). These vectors, termed pCOP88[sp-*tetM*], pCOP88[sp-*ermB*] (14), and pCOP88[sp-*vanB*], encode *cas9*, *cat*, and single guide-RNAs targeting either *tetM*, *ermB*, or *vanB*, respectively. The vectors were introduced into strains V583, OG1RF and fecal isolate 94-1 by electroporation. Because *E. faecalis* OG1RF does not encode any of the targeted genes, it served as a non-targeted control. As expected, we observed few to no transformants for the targeting constructs (Figure 5). Contrarily, nontargeting constructs were able to enter the cells (Figure 5). These data compare to that observed in Figure 4A, specifically for isolate 94-1 and further supports the notion of a sequence-specific impact on the entry of CRISPR-antimicrobial plasmids.

**Figure 5.**
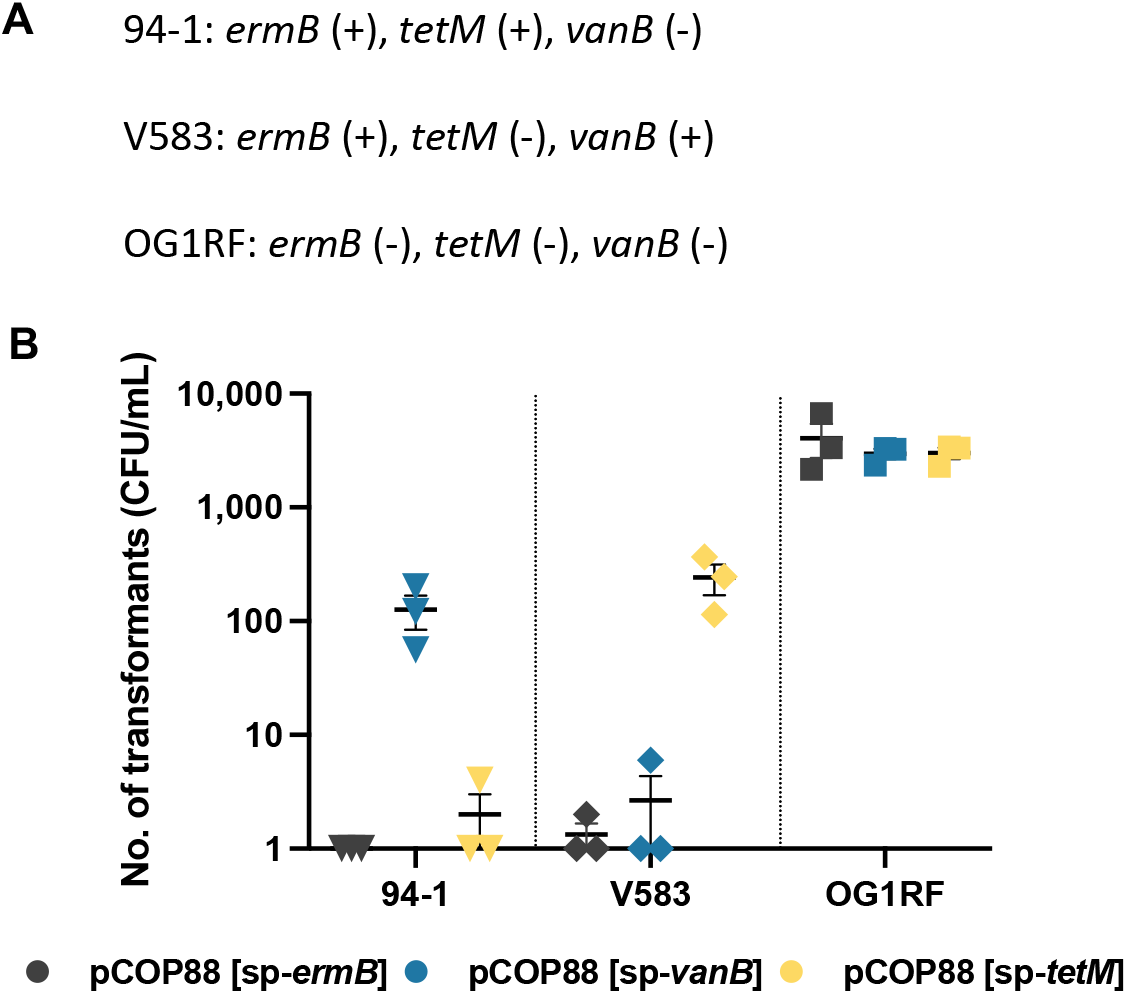
Sequence-specific effect on the entry of the pCOP88 vectors harboring the CRISPR-Cas machinery. (A) List of strains specifying the presence/absence of the genes targeted by the pCOP88 plasmids. (B) pCOP88 plasmids encoding cognate-RNA sequences yielded few to no transformants in strains 94-1 and V583, while plasmids encoding non-cognate guide-RNAs were able to enter the cells at higher rates. The OG1RF does not encode any of the targeted resistance genes. All three pCOP88 derivatives entered OG1RF similarly and at high rates. Error bars represent the geometric mean and SEM of data from three independent biological experiments.

### CRISPR-antimicrobial donors are antagonized by the cytolysin toxin

To investigate why the donors were being antagonized by some of the isolates, we used PCR to screen each isolate for the presence of cytolysin, a toxin with hemolytic and bacteriocin activity often found in *E. faecalis* (24, 33). Remarkably, we found that all isolates capable of antagonizing the donors harbor *cylA*, a gene present in the cytolysin operon (33) (Table 1). The complete cytolysin operon of all seven isolates was confirmed by whole genome sequencing analysis (Table S1). We further verified cytolysin activity in these isolates by detecting hemolysis on horse blood agar plates (Table 1).

Next, we assessed whether deletion of the cytolysin genes would improve donor survival when competing against the fecal isolates. We selected the cytolysin positive isolate 94-1, because of its low frequency of CRISPR-Cas antimicrobial plasmid acquisition (Figure 4A) and its ability to kill the donors (Figure 3). Deletion of the cytolysin genes from isolate 94-1 resulted in ∼4 to 5-log expansion of the donor population across both bacterial passages after conjugation (Figures S2 and S3), similar to that observed when conjugating with OG1SSp(pAM714) (Figure S2). These results show that the cytolysin toxin produced by *E. faecalis* fecal isolates is, at least partially, involved in the antagonization of the donor strain. Importantly, while better donor fitness resulted in a ∼3-log increase in transconjugant CFU/mL containing the nontargeting plasmid, it did not result in substantially higher conjugation of the targeting plasmids (Figure 4A) nor in the reduction of antibiotic resistance (Figure 2). Together, these results further demonstrate that there is a sequence-specific effect on conjugation of the CRISPR-Cas antimicrobial plasmids, which does not lead to sequence-specific removal of antibiotic resistance.

### Conjugation frequency is affected by factors other than poor donor fitness

Donor fitness is a key component for successful delivery and implementation of plasmid-based CRISPR-Cas antimicrobials. We therefore tested whether addition of the cytolysin immunity gene *cylI* (33) to the pKH88 plasmids would provide the donors with protection against the fecal isolates, thereby increasing conjugation frequency and improving depletion of antibiotic resistance. The modified CRISPR-Cas targeting plasmids were termed pDP01[sp-*tetM*] and pDP01[sp-*ermB*]. Unexpectedly, plasmid pDP01[sp-*vanB*] was found to have mutations in *cas9* upon routine plasmid sequence verification, therefore we utilized the nontargeting control plasmid pDP01[sp-NT]. The NT sequence is a random sequence that was confirmed by genome analysis to not occur in any of the recipient strains.

Conjugation assays between cytolysin-immune pDP01 donors and each cytolysin-positive isolate (Table 1) resulted in ∼2- to 6-log enhancement of donor survival compared to those harboring the pKH88 plasmid derivatives (Figure 3). However, similar to our previous observations, conjugation of the nontargeting plasmid (pDP01[sp-NT]) was detected in 6 of the 7 isolates at one or both passages, while transfer of either targeting plasmid (pDP01[sp-*ermB*] or pDP01[sp-*tetM*]), was only observed at passage 0, and near the limit of quantification (Figure 4B). In comparison, both targeting and nontargeting pDP01 derivatives conjugated into OG1SSp(pAM714) and OG1SSp(pCF10) at high frequencies (Figure 4B). These results show that donor fitness may not be the only factor responsible for the low conjugation frequency observed among these fecal isolates.

We hypothesized that one or more types of genome defense or anti-defense systems present in these fecal isolates may be preventing the successful delivery of the CRISPR-antimicrobial plasmids. We utilized Illumina and MinION sequencing to close the genomes of each fecal isolate selected for the study. We implemented genomic search strategies and the Prokaryotic Antiviral Defence LOCator (PADLOC) search database (34) to identify systems such as anti-CRISPR (35) and the antiplasmid defense system Wadjet (36), which could be responsible for the low conjugation frequency observed. We were unable to identify any sequences that matched the databases (Table S1). Our results warrant further investigation to determine the cause for the low conjugation observed among the fecal isolates.

### Bacteriocin 21 is required for the killing activity of the CRISPR-Cas antimicrobials

Despite the low conjugation frequency observed, fecal isolates 119-1, 119-2 and 152 had total erythromycin and tetracycline resistance reduction to near or at the level of detection when competing against the cytolysin-immune pDP01 donors (Figure 6). It is important to note that (i) in all three cases, resistance depletion also occurred when challenging the cells with the nontargeting plasmid and, (ii) addition of cytolysin immunity increased the donor survival considerably when competing against these isolates (Figure 3). Higher donor density in the population may have increased the concentration of Bac-21, produced by the donors, potentially killing susceptible recipients. The significant reduction of total resistance observed in these three fecal isolates was not observed when competing against the pKH88 derivatives, suggesting that these isolates may be sensitive to Bac-21 (Figure 7). Additionally, isolates displaying sensitivity towards the pKH88 antimicrobial derivatives (i.e., 59, 101-1, 133-1 and 142-1) were unable to kill the donors (Figure 2, Figure S2), further suggesting that the resistance reduction observed is due to a combination of CRISPR cleavage and the Bac-21 bacteriocin produced by the donors.

**Figure 6.**
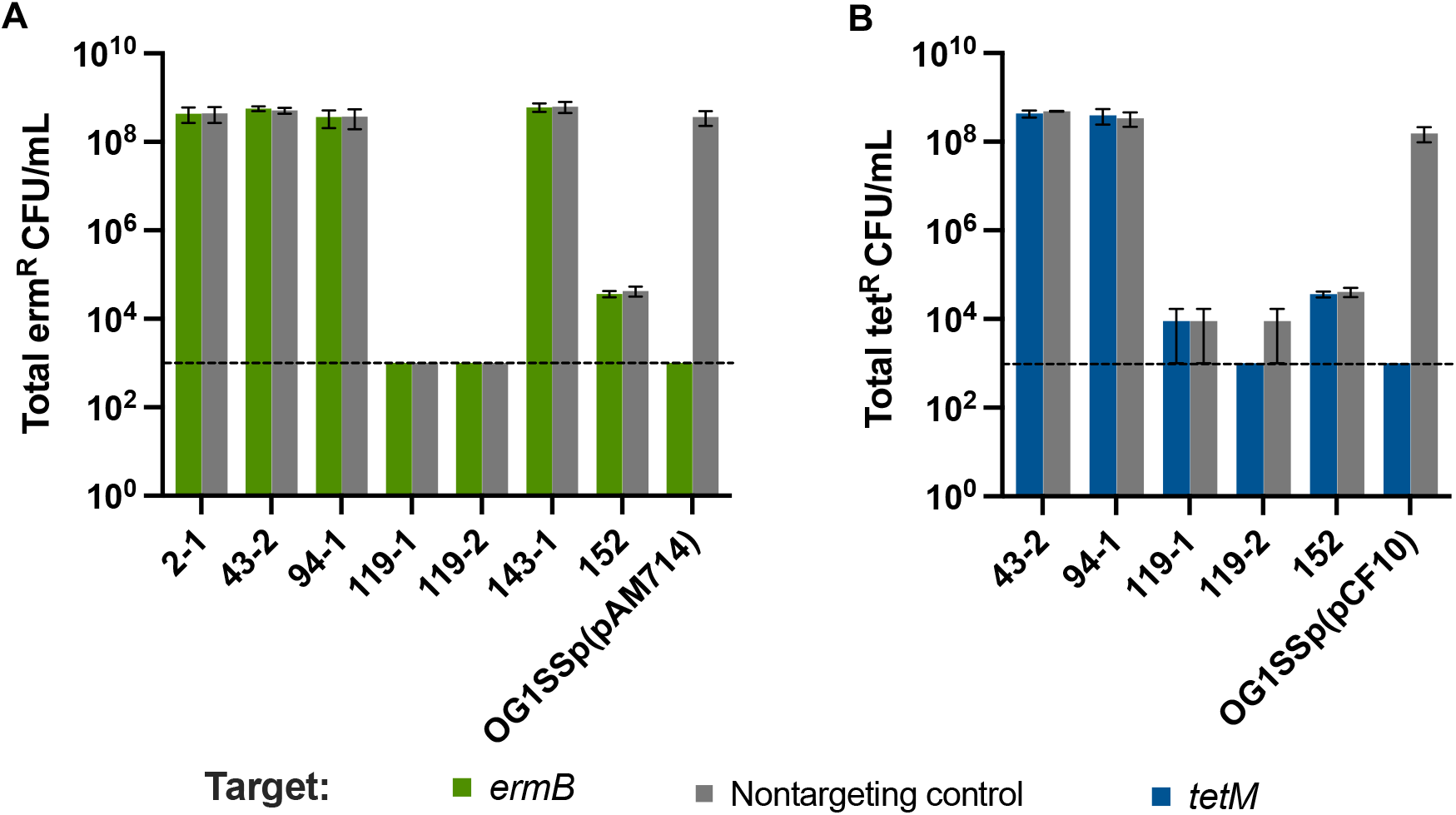
Antibiotic resistance depletion after treatment with pDP01 derivatives at passage 1. Sequence-specific resistance depletion of (A) erythromycin and (B) tetracycline was observed when targeting the *ermB* or *tetM* gene in the model strains OG1SSp(pAM714) and OG1SSp(pCF10), respectively. Fecal isolates 119-1, 119-2 and 152 had reduction of both erythromycin and tetracycline resistance, although not sequence-specific. Error bars represent the geometric mean and SEM of data from three independent biological experiments.

**Figure 7.**
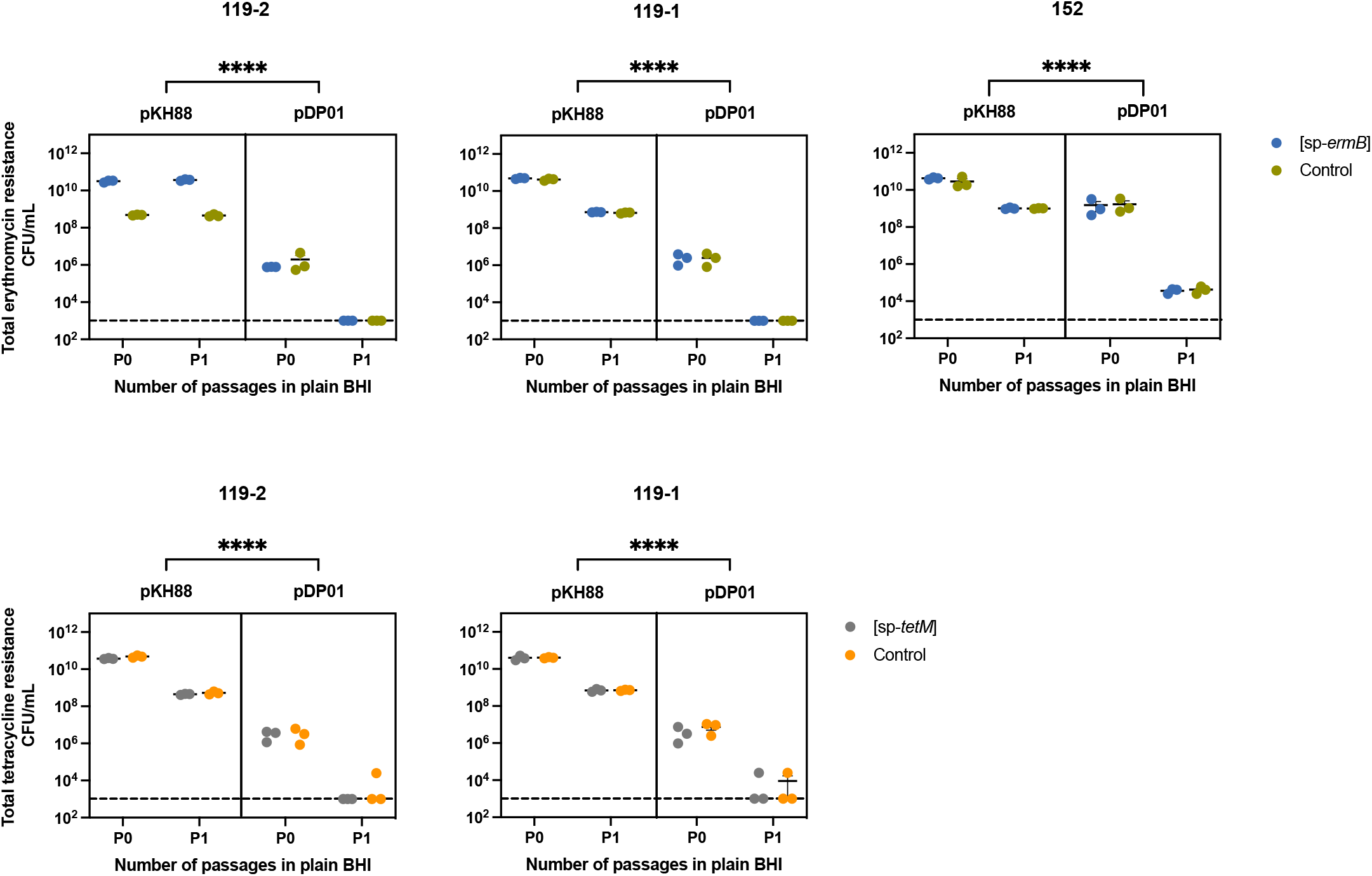
Total erythromycin and tetracycline resistance depletion after treatment with pKH88 versus pDP01. Isolates 59, 119-2 and 152 had a significant decrease of both antibiotic resistance phenotypes when treated with pDP01 compared to pKH88, although this reduction was not sequence-specific. Error bars represent the geometric mean and SEM of data from three independent biological experiments. Three-way ANOVA; ****, P= <0.0001.

To test this hypothesis, we deleted the gene *bacA*, encoding the bacteriocin precursor for Bac-21 (31), from pKH88[sp-*tetM*], generating pMA05[sp-*tetM*]. We compared the killing activity of donors with pMA05[sp-*tetM*] versus donors with pKH88[sp-*tetM*] against the model strain OG1SSp(pCF10) and the fecal isolate 59. Deletion of *bacA* resulted in a significant reduction in the killing activity of the donor against fecal isolate 59 (Figure 8A). Interestingly, without Bac-21 activity, the CRISPR-antimicrobial plasmid was unable to reduce tetracycline resistance in OG1SSp(pCF10) (Figure 8B). These results show that a critical component of tetracycline resistance reduction observed during CRISPR-Cas antimicrobial treatment is Bac-21-mediated cell death.

**Figure 8.**
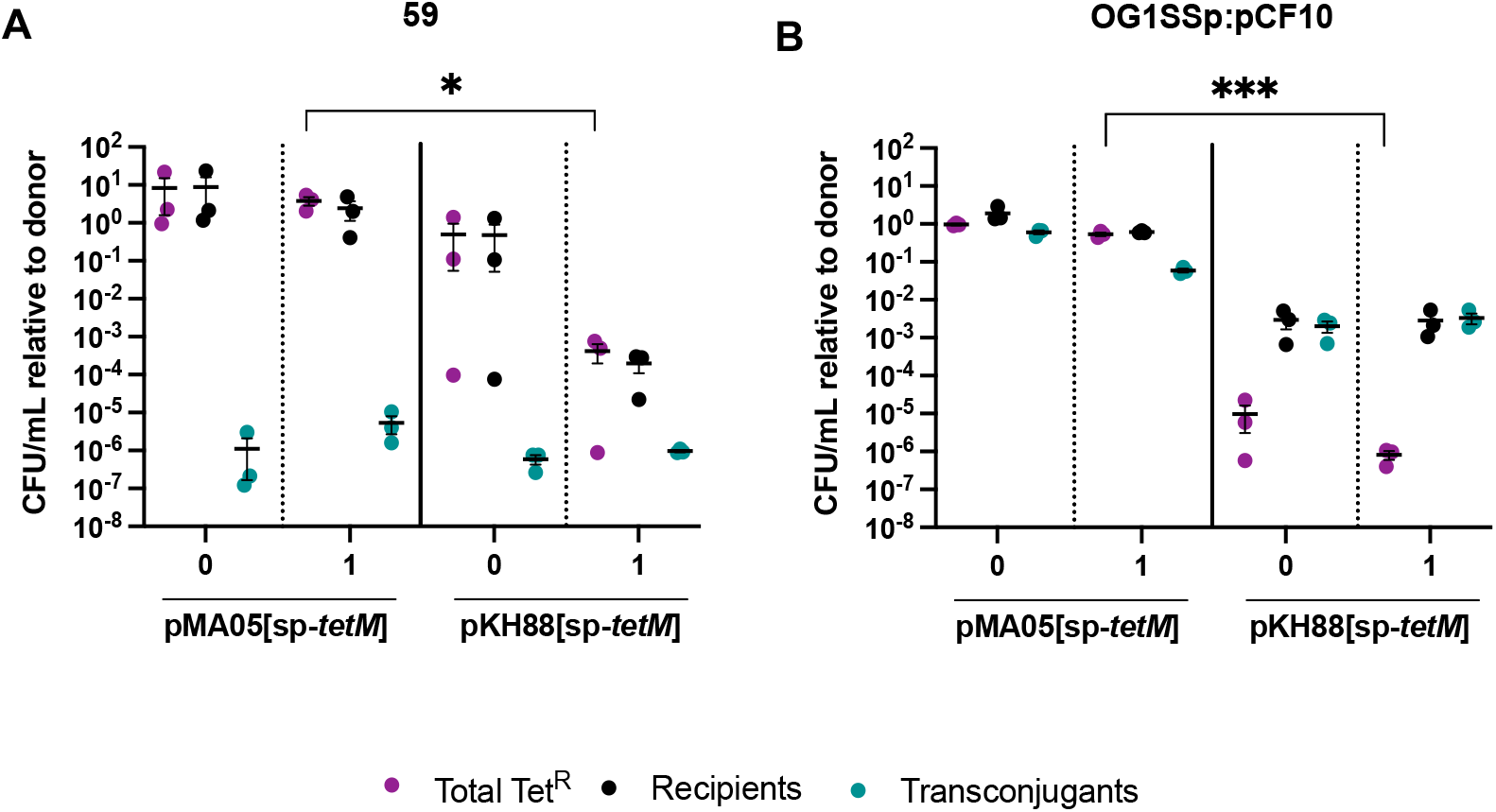
Loss of Bac21 activity reduces CRISPR-Cas antimicrobial killing activity. CRISPR-Cas antimicrobial lacking the Bac-21 precursor (pMA05[sp-*tetM*]) in the plasmid backbone has decreased targeting activity in A) fecal isolate 59 and B) OG1SSp(pCF10) compared to the wild type CRISPR-Cas antimicrobial plasmid (pKH88[sp-*tetM*]). Unpaired t-test; ***, P= 0.0006 (for OG1SSp(pCF10)); *, P= 0.0161 (for fecal isolate 59). Error bars represent the geometric mean and SEM of data from three independent biological experiments.

## DISCUSSION

In this study, we investigated the efficacy of our previously established CRISPR-Cas antimicrobials against a recent collection of fecal surveillance *E. faecalis* isolates. This collection represents ‘wild,’ uncharacterized bacteria, which are the intended targets for CRISPR-Cas antimicrobial therapy. Multiple groups —including our own, have demonstrated that CRISPR-Cas can be used as programmable sequence-specific antimicrobials (7, 10, 14, 21, 37, 38). The work cited here represents only a fraction of the many studies to date. Although these studies have established the groundwork for implementing CRISPR-Cas as an antimicrobial, most have focused primarily on examining CRISPR targeting in well-established model strains (11–14, 21). Therefore, we have limited knowledge of how effective CRISPR-based antimicrobials can be against uncharacterized, wild bacterial strains. Understanding potential limitations is important in order to develop a therapy that can achieve clinical application.

Our study employed *E. faecalis* donor strains to deliver CRISPR-Cas antimicrobial plasmids to either *E. faecalis* laboratory model strains or *E. faecalis* ‘wild’, uncharacterized fecal isolates. Our conjugation experiments revealed notable differences between these two groups: (i) Conjugations between the donor strain and most fecal isolates resulted in the antagonization of the donor. By contrast, competition with *E. faecalis* model strains (V583; OG1SSp) resulted in high donor survival. (ii) We show a remarkably higher plasmid acquisition by the model strains compared to the fecal isolates; this was especially true when using targeting constructs. (iii) Sequence-specific removal of antibiotic resistance was achieved in all model strains, consistent with our previous observations (14), while it was only partially attained in two of the fecal isolates.

The low survival of donor cells presumably indicates poor competitive ability against the wild fecal isolates compared to the model strains, possibly due to competitive factors encoded by these wild isolates. We note that our donor strain, CK135RF, and our model recipient, OG1SSp, are both derivatives of OG1, thus these strains are highly clonal; however, V583 is a MDR clinical isolate that is not clonal with OG1 and has a ∼0.6 Mbp larger genome (39, 40). OG1 was isolated in the 1970s (41), and V583 was isolated in the 1980s (30). Laboratory domestication of our model strains may have caused a loss of competitive fitness over time, rendering them less antagonistic against our donor strain. Multiple studies in various bacterial species have shown genetic differences upon laboratory domestication and storage, causing loss or changes of traits such as growth rates and virulence (25–28, 42). For instance, Müller et al. established that domesticated *Bacillus subtilis* strains had lost their ability to produce secondary metabolites conferring resistance against *Myxococcus xanthus* predation (42). In another example, a study performed in *Campylobacter jejuni* compared several of stocks of the same strain from different laboratories and found differences in their virulence (27).

Among the numerous competitive factors encoded by *E. faecalis* is cytolysin (24), a toxin with bactericidal activity found in highly virulent *E. faecalis* strains (22, 24). Indeed, we found that all seven fecal isolates that antagonized the donor encode this toxin, consistent with a previous *in vitro* study reporting that non-cytolytic enterococci strains were outcompeted by cytolytic strains (43). Unexpectedly, we show that improving donor fitness by incorporating cytolysin immunity did not directly translate to higher conjugation efficiencies among the cytolytic fecal isolates. In particular, acquisition of the CRISPR-Cas targeting plasmids remained low, suggesting the existence of other factors affecting conjugation. Because this phenotype was specific to the fecal isolates, we speculate that these isolates —being more recent, and less domesticated than our model strains— may be better adapted to prevent horizontal gene transfer of MGEs. Studies have shown that MGEs harbor many defense mechanisms (44). It has been argued that the high prevalence of defense systems in bacterial genomes is due to the continuous acquisition of MGEs harboring such systems (44). It is possible that these fecal isolates acquired multiple defense systems through HGT while existing in the gut microenvironment. Contrarily, it has been previously shown that laboratory evolution can cause the loss of defense systems (29), suggesting the possibility that our model strain may have gone through a similar evolutionary path during the many years of laboratory domestication. While we were unable to identify known anti-plasmid defense systems within the genomes of the fecal isolates, we cannot rule out the existence of such mechanisms within their genomes. The increasing number of defense systems found thus far across numerous bacterial genomes suggest that many of the genes with unknown functions may be defense systems (44).

However, this overall line of speculation may not explain why we observed a sequence-specific effect on the delivery of the CRISPR-Cas antimicrobial plasmids, i.e., all fecal isolates received the CRISPR-Cas antimicrobial plasmid with noncognate spacers, while the delivery of the plasmid with cognate spacers was unsuccessful. Consequently, we were unable to achieve sequence-specific reduction of antibiotic resistance among most of the fecal isolates. We also observed a sequence-specific effect on the plasmid entry into V583 in the transformation assays, however we were able to achieve plasmid delivery and sequence-specific depletion of resistance in V583 in the conjugation assays. The specific genetic differences between V583 and the fecal isolates that may explain these differing effects remain to be elucidated.

We observed that three of the fecal isolates were depleted from the population only when competing against the cytolysin-immune donors, regardless of the CRISPR-Cas plasmid being employed (cognate or noncognate spacer). Notably, this phenotype was lost in the absence of *bacA*, the precursor for the Bac-21 bacteriocin (31). It is plausible that improving donor survival directly increased the concentration of bacteriocin Bac-21, which has been previously shown to be secreted into the media (45), and that higher concentration of bacteriocin killed the sensitive isolates. Our results are consistent with a previous study demonstrating that Bac-21-positive *E. faecalis* display killing activity against other *E. faecalis* clinical isolates (45).

Our work revealed that our current delivery method for CRISPR-Cas antimicrobials is limited by the donor strain utilized. Moving forward, to implement conjugative plasmids as the delivery mechanism, we must improve the donor strain. Enhancements can be achieved by engineering an *E. faecalis* donor strain to be more competitive against wild strains. Another previously proposed avenue is to utilize gut-adapted bacteria, such as probiotic strains, which could be more resilient against other gut bacteria (46). Collectively, our results support our hypothesis that the differences observed between the two recipient groups are due to the presence (or lack thereof) of competitive factors. Although more studies are required to confirm our findings, our work emphasizes the need to study the efficacy of CRISPR-Cas antimicrobials in the context of wild bacterial strains to identify potential limitations. Understanding those limitations will help us develop delivery mechanisms adept to endure obstacles found within the gut microbiota and delivery the CRISPR-Cas load successfully, which is key for a viable therapeutic.

## MATERIALS AND METHODS

### Bacterial strains, growth conditions and routine molecular biology techniques

All bacterial strains and plasmids used in this study are listed in Tables S2 and S3, respectively. *Escherichia coli* strains EC1000 and Top10, used for plasmid propagation, were cultured in Lysogeny Broth (LB) at 37°C with agitation at 220 rpm. *E. faecalis* strains were grown in Brain-Heart Infusion (BHI) medium without shaking at 37°C, unless otherwise specified. When needed, antibiotics were applied for *E. faecalis* and *E. coli* in the following concentrations: chloramphenicol, 15 µg/mL; erythromycin, 50 µg/mL; tetracycline, 10 µg/mL; vancomycin, 10 µg/mL; rifampicin, 50 µg/mL; fusidic acid, 25 µg/mL; spectinomycin, 500 µg/mL; streptomycin, 500 µg/mL. Electrocompetent *E. coli* and *E. faecalis* cells were prepared as previously described (16). Routine PCR analysis was conducted with *Taq* DNA polymerase (New England Biolabs). PCR for cloning purposes was performed with Q5 DNA polymerase (New England Biolabs). DNA oligo primers were obtained from Sigma-Aldrich (Table S4). PCR products were purified using the PureLink PCR purification kit (Invitrogen) per manufacturer protocols. Plasmids were isolated with the GeneJet plasmid miniprep purification kit (Thermo Fisher) per manufacturer protocols. Routine Sanger DNA sequencing was carried out at the Massachusetts General Hospital DNA Core Facility. Genomic DNA was extracted using DNeasy Blood and Tissue kit (Qiagen) according to the manufacturer’s instructions with a few modifications: prior to using the kit, the cells were pretreated with 180 μL of lysozyme (50 mg/mL), 15 μL of preboiled RNase A (20 mg/mL) and 25 μL of mutanolysin (2,500 U/mL) and incubated at 37°C for 2 h. All plasmids generated were assembled using Gibson Assembly Master Mix (New England Biolabs) per manufacturer protocols. All primers utilized in this study are found in table S4.

### Strain and plasmid gene deletions

Plasmids used to genetically manipulate *E. faecalis* were originated from derivatives of plasmid pLT06 (47). The resulting derivatives, termed pCOP88, previously described in Rodrigues et al. 2019, maintained chloramphenicol selection, p-Cl-phenylalanine counterselection and temperature sensitive allelic-exchange qualities. To generate *E. faecalis* 94-1Δ*cyl*, pCOP fragments encoding genes *pheS*, *oriT* (with temperature sensitive *repA*) and *cat*, and 1.0 kb up and downstream sequences of the cytolysin genes in *E. faecalis* 94-1 were amplified and assembled as described above, creating plasmid pKO1 (Figure S4). To generate the CRISPR-Cas antimicrobial pMA05 derivatives, pCOP fragments encoding genes *pheS*, *oriT* and *cat*, and 1.0 kb up and downstream sequences of the *bacA* gene in plasmid pPD1 were amplified and assembled, creating plasmid pCOP212. Next, 1 μg of the resulting plasmid was transformed into electrocompetent *E. faecalis* 94-1 and *E. faecalis* CK135RF(pPD1) respectively, and incubated at the permissive temperature of 30°C for 48 h. Following 30°C incubation, cells were streaked on a fresh BHI plate and shifted to 42°C. After 48 h of incubation, individual colonies were inoculated in 1 mL of fresh BHI and incubated at 37°C for 5 h, followed by plating on MM9YEG supplemented with p-chloro-phenylalanine. To confirm cytolysin deletion in strain 94-1, recombinants were screened by PCR and analyzed for loss of hemolysis activity, as described below. To confirm deletion of *bacA,* CK135RF(pPD1Δ*bacA*) was spotted onto a bacterial lawn susceptible to Bac-21. Next, previously described integration plasmid pCOP88[sp-*tetM*] (14) was transformed into electrocompetent CK135RF(pPD1Δ*bacA*) and recombination was performed as described above. The resulting pPD1 plasmid lacked the *bacA* gene while harboring the CRISPR-Cas antimicrobial machinery (14). The plasmid was termed pMA05[sp-*tetM*]. pCOP88 plasmids insert the engineered region P*_bacA_*-*cas9*-*cat*-guide RNA into the plasmid pPD1 between ORFs ppd4 and ppd5.

### Construction of non-cytolysin immune CRISPR-Cas targeting vector

To generate a pKH88 derivative targeting gene *vanB*, integration plasmid pCOP88[sp-*tetM*] (14) was linearized, excluding its CRISPR array. Then, the CRISPR array of plasmid pGR-*vanB* was amplified and assembled with the linearized pCOP88 as mentioned above (Figure S5A). To generate a pCOP88 derivative with a nontargeting spacer, previously engineered pCOP88[sp-*tetM*] plasmid was linearized excluding the spacer. Amplification was performed using primers containing overhangs encoding for a random nontargeting sequence (NT) (Figure S5B). Plasmids were assembled using the NEBuilder HiFi DNA Assembly. Next, the newly assembled plasmids pCOP88[sp-*vanB*] and pCOP88[sp-NT], respectively, were transformed into electrocompetent *E. faecalis* CK135RF harboring the native pPD1 plasmid. Temperature shifts and MM9YEG plating were implemented as previously described. Recombination resulted in the generation of plasmids pKH88[sp-*vanB*] and pKH88[sp-NT], respectively.

### Construction of cytolysin immune CRISPR-Cas targeting vector

To generate pPD1-*cylI*, the *cylI* coding sequence from *E. faecalis* OG1SSp(pAM714) was subcloned with the P_rpsL_ promoter and amplified as one fragment. PCR product P_rpsL_-*cylI* was assembled as described above, with 1.0 kb sequences up and downstream of the chosen insertion site along with pCOP88 fragments (encoding genes as described above), generating plasmid pCOP214 (Figure S6A). The resulting plasmid was transformed into electrocompetent *E. faecalis* CK135RF:pPD1. To obtain recombinants, transformed cells underwent the same procedure as described above. P_rpsL-_*cylI* was inserted between open reading frames (ORFs) ppd20 and *nrdH* gene. To confirm gene insertion, recombinants were screened by PCR and assessed for acquisition of cytolysin immunity as follows: by spotting 10 μL of OG1SSp(pAM714) (a cytolytic strain) onto individual lawns of bacteria presumed to be recombinants. Recombinants not showing a zone of inhibition were selected for further manipulation. Once pPD1-*cylI* was established, electrocompetent CK135RF(pPD1-*cylI*) cells were transformed with either pCOP88[sp-*tetM*], pCOP88[sp-*ermB*], pCOP88[sp-*vanB*] or pCOP88[sp-NT]. Integration resulted in the generation of plasmids pDP01[sp-*tetM*], pDP01[sp-*ermB*], pDP01[sp-*vanB*] and pDP01[sp-NT] respectively (Figure S6B). The sequence integrity of all the CRISPR-Cas antimicrobial plasmid derivatives was confirmed by whole genome sequencing using the Illumina platform at the Microbial Genome Sequencing Center (MiGS).

### Conjugation assays

*E. faecalis* donors and recipients were individually cultured in BHI with appropriate antibiotic selection overnight. The next day, cultures were diluted to 0.1 OD_600nm_ into fresh BHI and incubated at 37°C for 1.5 h until exponential phase was reached. A mixture of 900 μL of recipient and 100 μL of donor was pelleted and plated on BHI medium without antibiotics. After overnight incubation at 37°C, the cell lawns were scraped into 2 mL 1X PBS supplemented with 10 mM EDTA. Serial dilutions were plated on BHI plates supplemented with antibiotics to select for donors, recipients, transconjugants and total antibiotic resistance (Passage 0). For conjugation passages, the recovered bacterial mixture was diluted 1:10,000 in fresh BHI without selection and incubated at 37°C overnight. After this, the cultures were serially diluted and plated as described above (Passage 1). To account for potential spontaneous streptomycin resistance mutations in the donor population, 100 μL of donor monocultures were plated on BHI plates supplemented with streptomycin and grown overnight at 37°C. Colonies were counted and CFU/mL was determined.

### Pheromone induction assays

Pheromone induction was performed as previously described with some modifications (48). Overnight cultures were diluted 1:10 in fresh BHI containing no antibiotic and were incubated statically 1.5 h at 37°C in the presence of synthetic pheromone cPD1 (GL Biochem, Shanghai) at a concentration of 0.5 μg/mL. Cell clumping was assessed by visual observation. Assays were carried out in biological triplicates.

### Screening of fecal surveillance isolates

All *E. faecalis* fecal isolates (19) were screened by colony PCR for the genes *ermB, tetM* and *cylA* using primer pairs listed in Table S4. Presence of those genes was reconfirmed by manually searching for their sequences in complete genome sequences of the fecal isolates. Strains were tested for antibiotic resistance phenotypes by streaking onto BHI agar plates supplemented with antibiotics of interest and examining bacterial growth after 24 h incubation at 37°C (Table 2). *vanB* screening was previously performed (19).

### Illumina sequencing

DNA was sequenced at the Microbial Genome Sequencing Center (MiGS), where samples were processed to the 300 Mbp depth on the NextSeq 2000 platform, or at the Genome Center at the University of Texas at Dallas, where sample libraries were pooled and sequenced with a paired-end read of 2 × 150 bp on the NextSeq 500 sequencing platform.

### MinION sequencing

Fecal isolates where sequenced by Oxford Nanopore technology (ONT) (49) as previously described (50). Briefly, a total of 1.5 µg of genomic DNA was mechanically sheared into 8 kb fragments using Covaris g-tubes per manufacturer’s instructions prior to preparing the library using the ONT Ligation Sequencing Kit 1D (SQK-LSK108). Multiplexed samples were sequenced using the Native Barcoding Kit (EXP-NBD103) in each flowcell. Libraries were base-called with MinKNOW (v3.5.5) to generate fastq and fast5 sequence reads.

### Hybrid genome assembly of the *E. faecalis* fecal isolates

Assembly of the fecal isolate genomes was performed by employing a combination of MinION sequencing reads and Illumina sequencing short reads as previously described (51). Genome assemblies were annotated with PROKKA (v 1.12) (52). Complete genome sequences of all *E. faecalis* isolates used in this study have been deposited in the NCBI database under the following accession numbers: CP091906-CP091907, CP091904-CP091905, CP091901-CP091903, CP091908-CP091911, CP091899-CP091900, CP091897-CP091898, CP091895-CP091896, CP091892-CP091894, CP091889-CP091891, CP091886-CP091888 and CP091884-CP091885.

### Genome analysis of the fecal isolates

Multi-locus sequence typing (MLST) was determined using the *E. faecalis* MLST database (http://pubmlst.org/efaecalis/) (53). Antibiotic resistance genes were found using ResFinder 4.0 (54). To identify CRISPR loci and *cas* genes the web tool CRISPRCasFinder was utilized with manual verification (55, 56). Searches for anti-plasmid defense systems anti-CRISPR and Wadjet were done using the AcrDB (57) and PADLOC (34) respectively. Bacteriocins were identified using the BAGEL4 web server (58) (Table S1). The cytolysin operon of each positive isolate was manually inspected to confirm the operon was complete.

### Hemolysis assay

Qualitative detection of hemolysis activity was performed using BHI agar plates supplemented with 5% (v/v) horse blood (Fisher Scientific). 10 μL of overnight culture were spotted on blood plates and incubated at 37°C. Following 24 h incubation, the plates were assessed for zones of hemolysis. Assays were carried out in biological triplicates.

### Generation of streptomycin-resistant mutants

Streptomycin-resistant derivatives of *E. faecalis* fecal isolates were generated by selecting for spontaneous mutants after plating 100 μL of cultured bacteria on BHI supplemented with 500 μg/mL streptomycin and incubating at 37°C for 24 h. Once colonies were obtained, three-day passages in broth BHI were performed. After the passages, cultures were streaked on BHI plates supplemented with 500 μg/mL streptomycin to confirm the stability of resistance.

### *E. faecalis* fecal isolate transformations

Electrocompetent cells were prepared as previously described (16) with minor modifications. Cells were grown overnight at 37°C in BHI. The next day, cultures were diluted 1:10 and incubated until the cells reached an OD_600nm_ of 0.5-0.75. Cells were pelleted by centrifugation and washed once with 10% glycerol, followed by treatment with 30 μg/mL lysozyme solution and incubation for 20 min at 37°C. Cells were then washed in ice-cold electroporation solution four times and resuspended in 300 μL electroporation solution. 50 μL of the resuspended cells were transformed with 1 μg of either pCOP88[sp-*ermB*], pCOP88[sp-*tetM*], or pCOP88[sp-*vanB*], using an Eporator (Eppendorf) and recovered in 1 mL of BHI supplemented with 0.5 M sucrose for 3 h at 30°C. After the recovery time, the cells were spun down and resuspended in 200 µL of fresh BHI. The resuspended cells were divided and plated as follows: 100 µL were directly plated on BHI plates supplemented with chloramphenicol. The remaining 100 µL were serially diluted, and 100 µL of each dilution was plated on BHI or BHI supplemented with chloramphenicol. Plates were incubated at 30°C. After an incubation period of 24-48 h, colonies were counted, and CFU/mL was calculated. To avoid variability in transformation efficiency, the same batch of prepared competent cells was transformed with each plasmid.

### Statistical and data analysis

Due to the high variation in CFU of donors obtained and that in some instances the recipient population decreased upon treatment with the CRISPR-Cas antimicrobials, the data were not normalized to transconjugant/donor as is commonly utilized in conjugation experiments. Instead, the data are presented directly as CFU/mL, except data specific to strain V583 in which case the CFU/mL were normalized to the donors. When 0-1 colonies were counted on a plate, a CFU/mL threshold was determined assuming one single colony was observed. Based on this, the limit of detection was established to be 1,000 CFU/mL. When 2-25 colonies were observed, the CFU/mL was calculated by assuming 25 colonies were counted. Based on this, the limit of quantification was established to be 25,000 CFU/mL.

GraphPad Prism (Version 9.1.1) was used to determine statistical significance. Depending on the experimental set up, the following statistical analysis were performed. For transformation efficiency, the homogeneity of variance was first established using the Levene’s test using R (R 3.3.2 GUI 1.68 Mavericks build (7288), then an unpaired Student *t* test assuming log-normal distribution was performed. For antibiotic depletion of targeted strains relative to nontargeted strains, multiple unpaired Student (Welch) *t* tests were performed, and for V583 analysis a Mann-Whitney test was performed. All assuming log-normal distribution. For differences between groups treated with plasmids pKH88 and pDP01, a 3-way ANOVA was performed.

## ACKNOWLEDGMENTS

This work was supported by R01AI116610 and the Cecil H. and Ida Green Chair in Systems Biology Science to KLP. We are grateful to Matthew Johnson for discussions about bioinformatic identification of anti-CRISPR systems.

## Supplemental figures and tables

**Figure S1.**
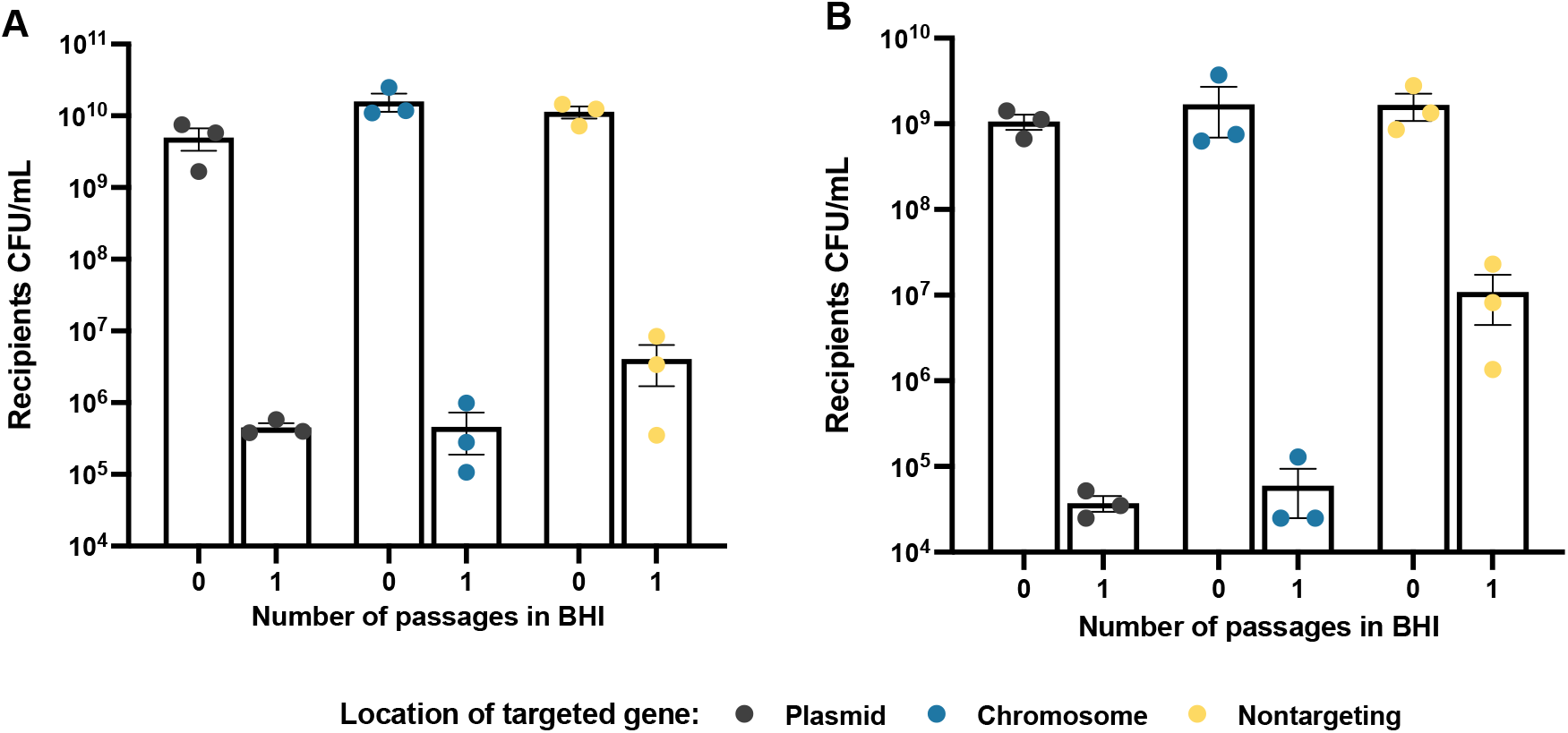
Reduction in recipient cell numbers upon CRISPR-Cas antimicrobial treatment is independent of target location. Fecal isolates (A) 59 and (B) 101-1 had sequence-specific reduction of viable cells regardless of whether the target was located on a plasmid or the chromosome relative to the nontargeting control. Threshold for sequence-specific reduction was established at 10-fold change or higher. Error bars represent the geometric mean and SEM of data from three independent biological experiments.

**Figure S2.**
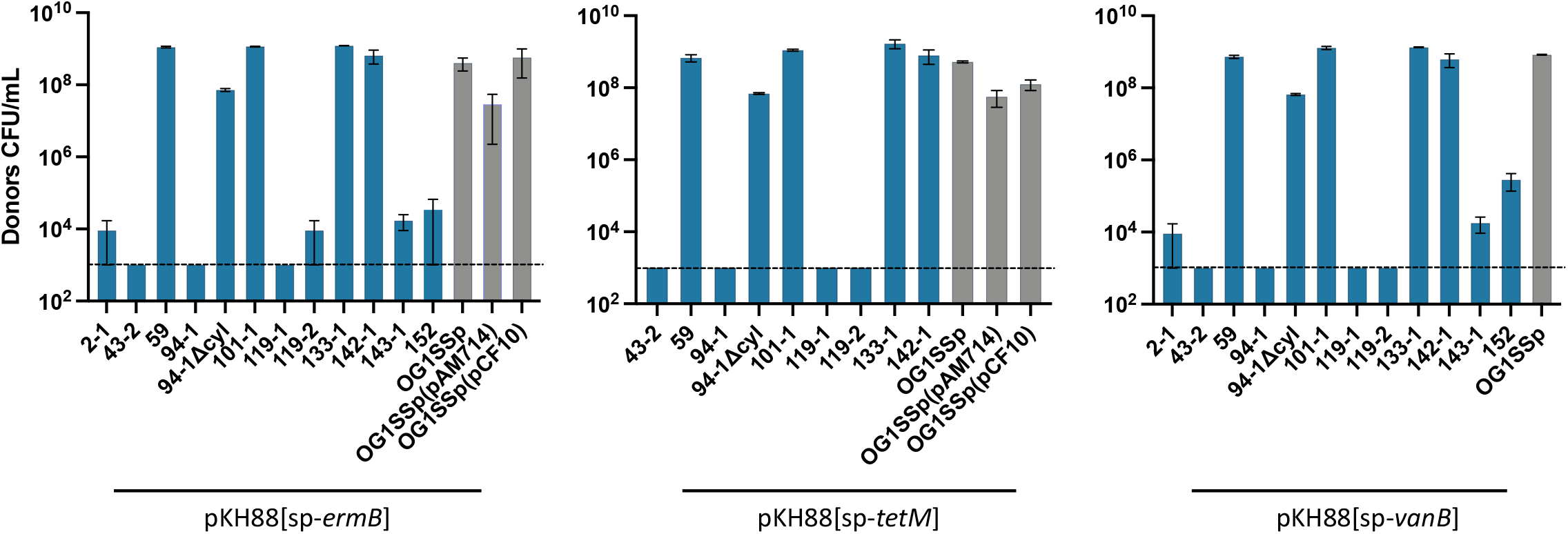
Donors of pKH88-based CRISPR-Cas antimicrobials compete poorly against some fecal isolates. Regardless of the pKH88 derivative, the population of donors fell either below the LOQ (2.5×10^4^) or LOD (1×10^3^) by the second passage when in presence of 7 (out of 11) fecal isolates. Blue bars represent fecal isolates, grey bars represent common *E. faecalis* model strains. Notably, fecal isolate 94-1 lacking the cytolysin genes (94-1Δcyl) increased donor CFU/mL by 4 to 5-log compared to the wild type 94-1. Error bars represent the geometric mean and SEM of data from three independent biological experiments.

**Figure S3.**
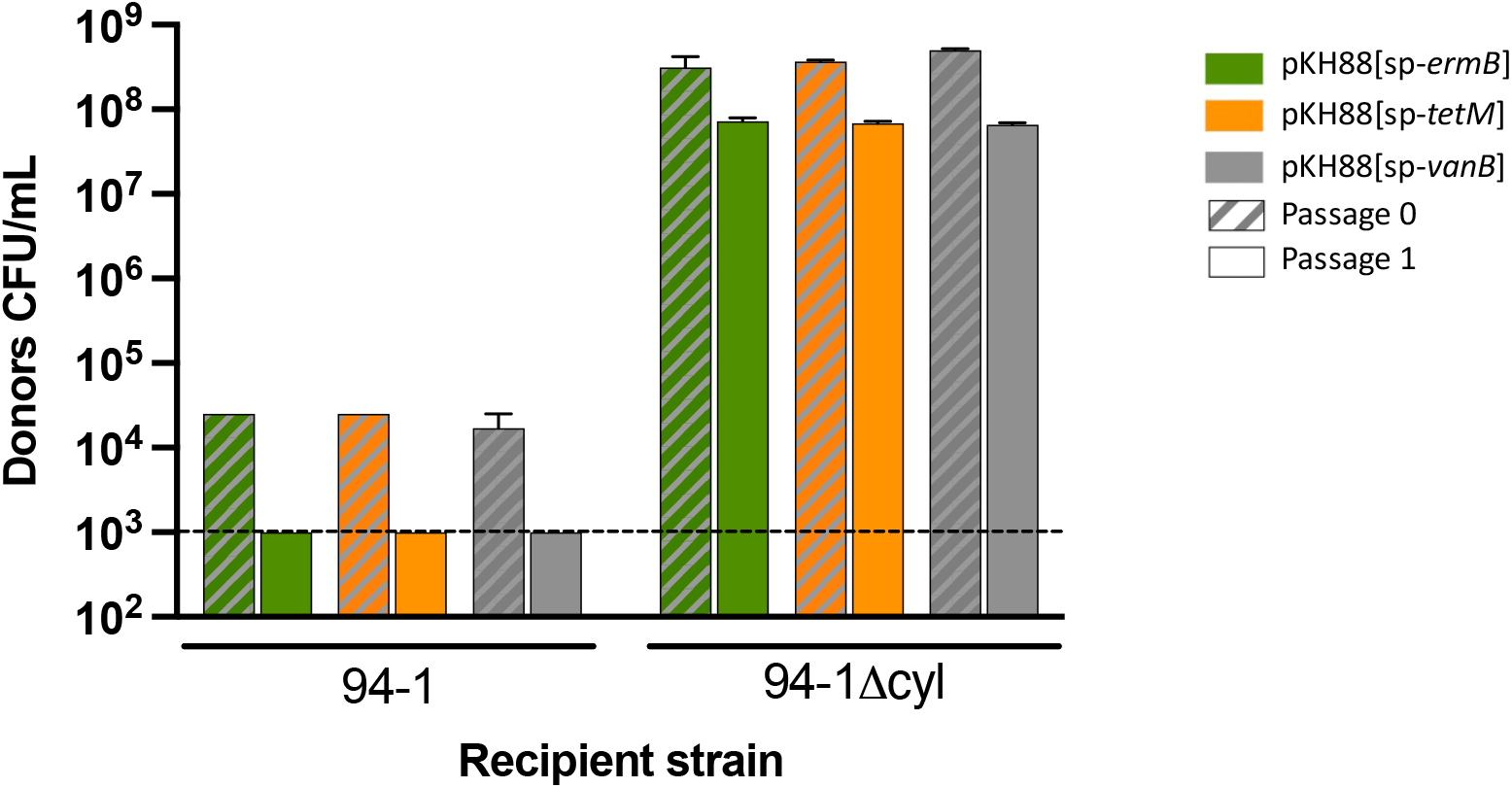
Deletion of the cytolysin operon in isolate 94-1 increases pKH88 donor survival. Donors competed better when fecal isolate 94-1 did not encode the cytolysin toxin, reflected by a 4 to 5-log increase in donor CFU/mL. This trend was maintained throughout both passages. Error bars represent the geometric mean and SEM of data from three independent biological experiments.

**Figure S4.**
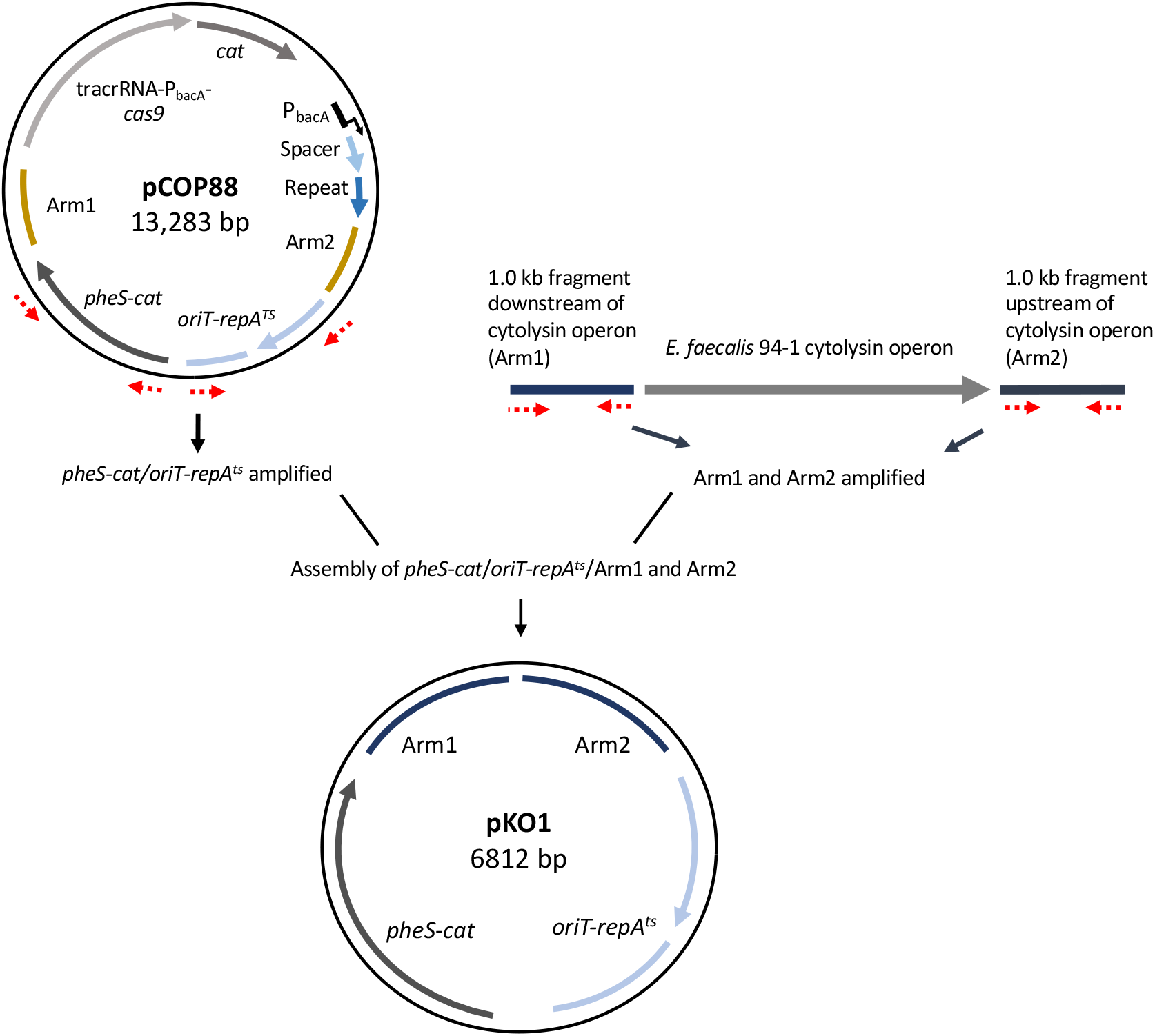
Strategy for the generation of plasmid pKO1, used to delete the cytolysin operon from isolate 94-1. pKO1 derivative encodes parts of the plasmid pCOP88, including the counter-selectable marker *pheS* and the temperature sensitive *repA* (*repA^ts^)* that allows integration into the chosen integration site at the non-permissive temperature of 42°C. 1-kb homology sites flanking the cytolysin operon in isolate 94-1 were cloned into pKO1 to facilitate homologous recombination and deletion of the operon.

**Figure S5.**
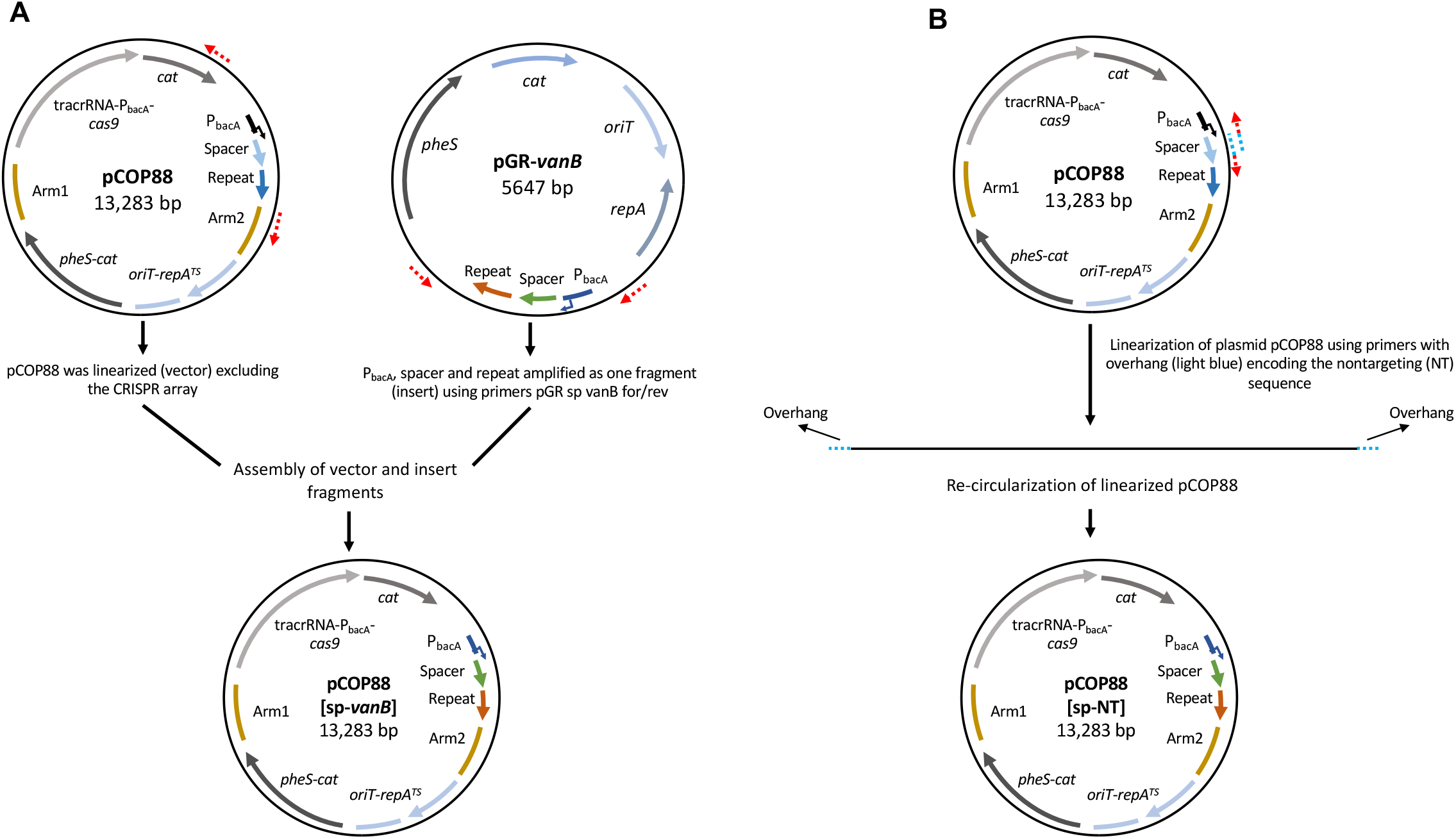
Strategy for the creation of the plasmid vectors pCOP88[sp-*vanB*] and pCOP88[sp-NT]. (A) Previously engineered pCOP88 plasmid was linearized excluding the sequence encoding the CRISPR array, while the CRISPR array encoding a *vanB* spacer was amplified from plasmid pGR-*vanB*. Fragments were assembled using the NEBuilder HiFi DNA Assembly, resulted in pCOP88[sp-*vanB*]. Dotted red arrows indicate the site for primer annealing and extension direction. (B) Previously engineered pCOP88 plasmid was linearized excluding only the already existing spacer and was replaced using primers designed with overhangs encoding the nontargeting (random) spacer sequence NT. The linearized product that included the new NT spacer was then re-circularized using the NEBuilder HiFi DNA Assembly, resulted in pCOP88[sp-NT]. Dotted red arrows indicate the site for primer annealing and extension direction, dotted blue indicates the overhang sequence of the primer.

**Figure S6.**
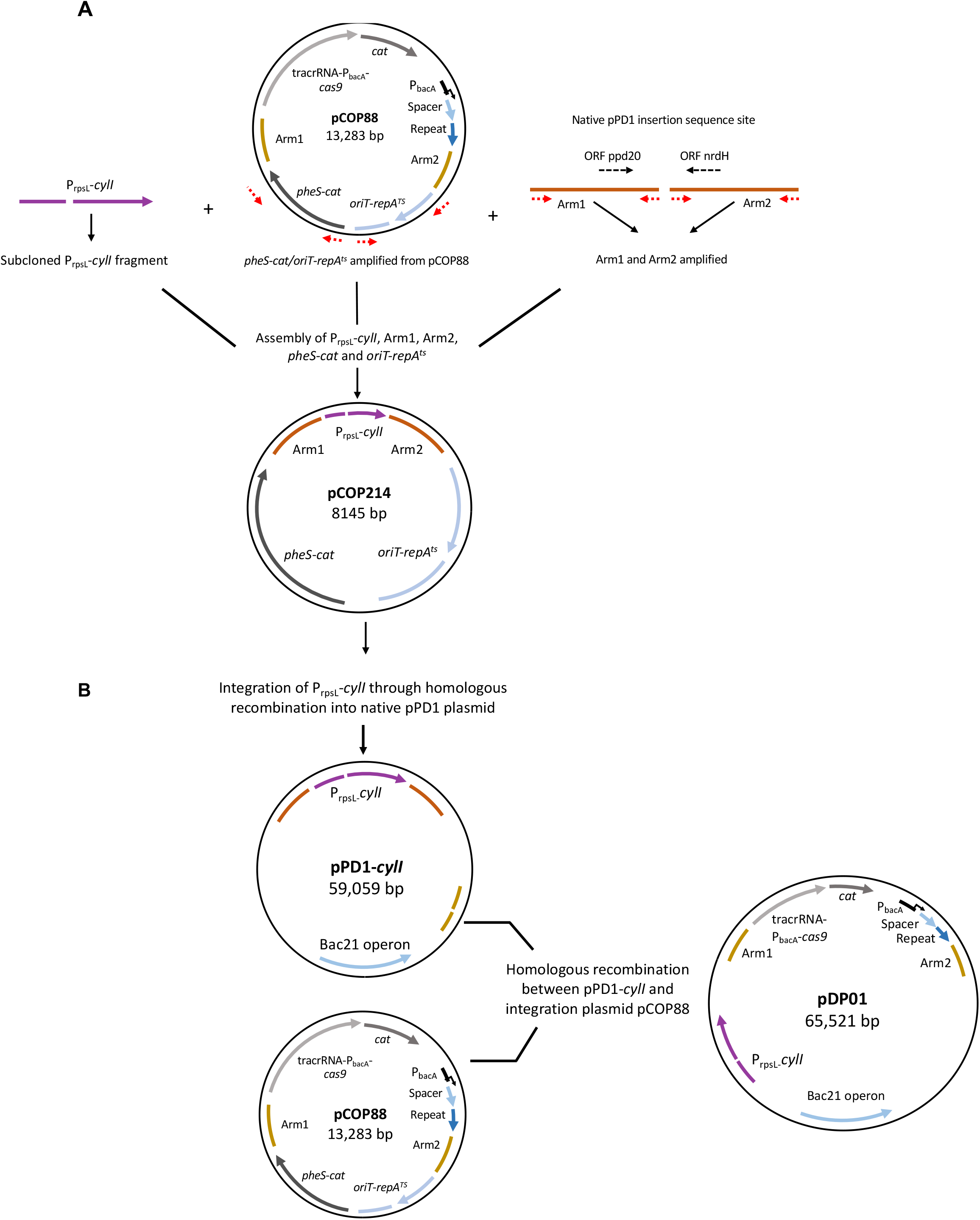
Strategy for construction of CRISPR-antimicrobial plasmid pDP01. (A) *cylI* coding sequence from *E. faecalis* OG1SSp(pAM714) was subcloned with the P_rpsL_ promoter, amplified as one fragment and assembled with 1.0 kb sequences up and downstream of the pPD1 insertion site to produce pCOP214. (B) pCOP214 was transformed into CK135RF(pPD1). Recombination was obtained by allelic exchange of the homologous sequences, yielding CK135RF(pPD1-*cylI*). Finally, those recombinants were transformed with different pCOP88 derivatives (pCOP88[sp-*ermB*], pCOP88[sp-*tetM*], pCOP88[sp-*vanB*]) or pCOP88[sp-NT]. Integration of the engineered region P_bacA_-*cas9*-*cat*-guide RNA resulted in the generation of plasmids pDP01[sp-*ermB*], pDP01[sp-*tetM*], pDP01[sp-*vanB*] and pDP01[sp-NT] respectively.

**Table S2.** Strains used in this study.

| Organism | Strain name | Description | Reference |
| --- | --- | --- | --- |
| *E. coli* | EC1000 | Cloning host, providing *repA* in trans. F- , araD139 (*araABC-leu*)7679, *galU*, *galK*, *lacX74*, *rspL*, *thi*, *repA* of pWV01 in *glgB*, km | 1 |
| *E. coli* | Top10 | F-mcrA Δ(*mrr-hsdRMS-mcrBC*) φ80lacZΔM15 ΔlacX74 nupG recA1 araD139 Δ(*ara-leu*)7697 galE15 galK16 rpsL(StrR) endA1 λ- | Invitrogen |
| *E. faecalis* | OG1SSp | Spectinomycin and streptomycin-resistant derivative of human oral cavity isolate OG1 | 2 |
| *E. faecalis* | OG1RF | Rifampin and fusidic acid-resistant derivative of human oral cavity isolate OG1 | 2 |
| *E. faecalis* | CK135RF | Spontaneous fusidic acid-resistant derivative of CK135 | 3 |
| *E. faecalis* | V583 | MDR bloodstream isolate, resistant to vancomycin, gentamicin, and erythromycin | 4 |
| *E. faecalis* | 2-1 | Fecal surveillance isolate. Collected in 2015 | 5 |
| *E. faecalis* | 43-2 | Fecal surveillance isolate. Collected in 2015 | 5 |
| *E. faecalis* | 59 | Fecal surveillance isolate. Collected in 2015 | 5 |
| *E. faecalis* | 94-1 | Fecal surveillance isolate. Collected in 2015 | 5 |
| *E. faecalis* | 101-1 | Fecal surveillance isolate. Collected in 2015 | 5 |
| *E. faecalis* | 119-1 | Fecal surveillance isolate. Collected in 2015 | 5 |
| *E. faecalis* | 119-2 | Fecal surveillance isolate. Collected in 2015 | 5 |
| *E. faecalis* | 133-1 | Fecal surveillance isolate. Collected in 2015 | 5 |
| *E. faecalis* | 142-1 | Fecal surveillance isolate. Collected in 2015 | 5 |
| *E. faecalis* | 143-1 | Fecal surveillance isolate. Collected in 2015 | 5 |
| *E. faecalis* | 152 | Fecal surveillance isolate. Collected in 2015 | 5 |
| *E. faecalis* | 94-1Δcyl | 94-1 fecal isolate derivative with an in-frame deletion of the cytolysin operon | This study |
| *E. faecalis* | 94-1SΔcyl | 94-1 fecal isolate derivative with an in-frame deletion of the cytolysin operon; Streptomycin-resistant | This study |
| *E. faecalis* | 2-1S | 2-1 fecal isolate derivative with streptomycin resistance | This study |
| *E. faecalis* | 59S | 59 fecal isolate derivative with streptomycin resistance | This study |
| *E. faecalis* | 94-1S | 94-1 fecal isolate derivative with streptomycin resistance | This study |
| *E. faecalis* | 119-1S | 119-1 fecal isolate derivative with streptomycin resistance | This study |
| *E. faecalis* | 119-2S | 119-2 fecal isolate derivative with streptomycin resistance | This study |
| *E. faecalis* | 142-1S | 142-1 fecal isolate derivative with streptomycin resistance | This study |
| *E. faecalis* | 143-1S | 143-1 fecal isolate derivative with streptomycin resistance | This study |
| *E. faecalis* | 152S | 152 fecal isolate derivative with streptomycin resistance | This study |

**Table S3.**
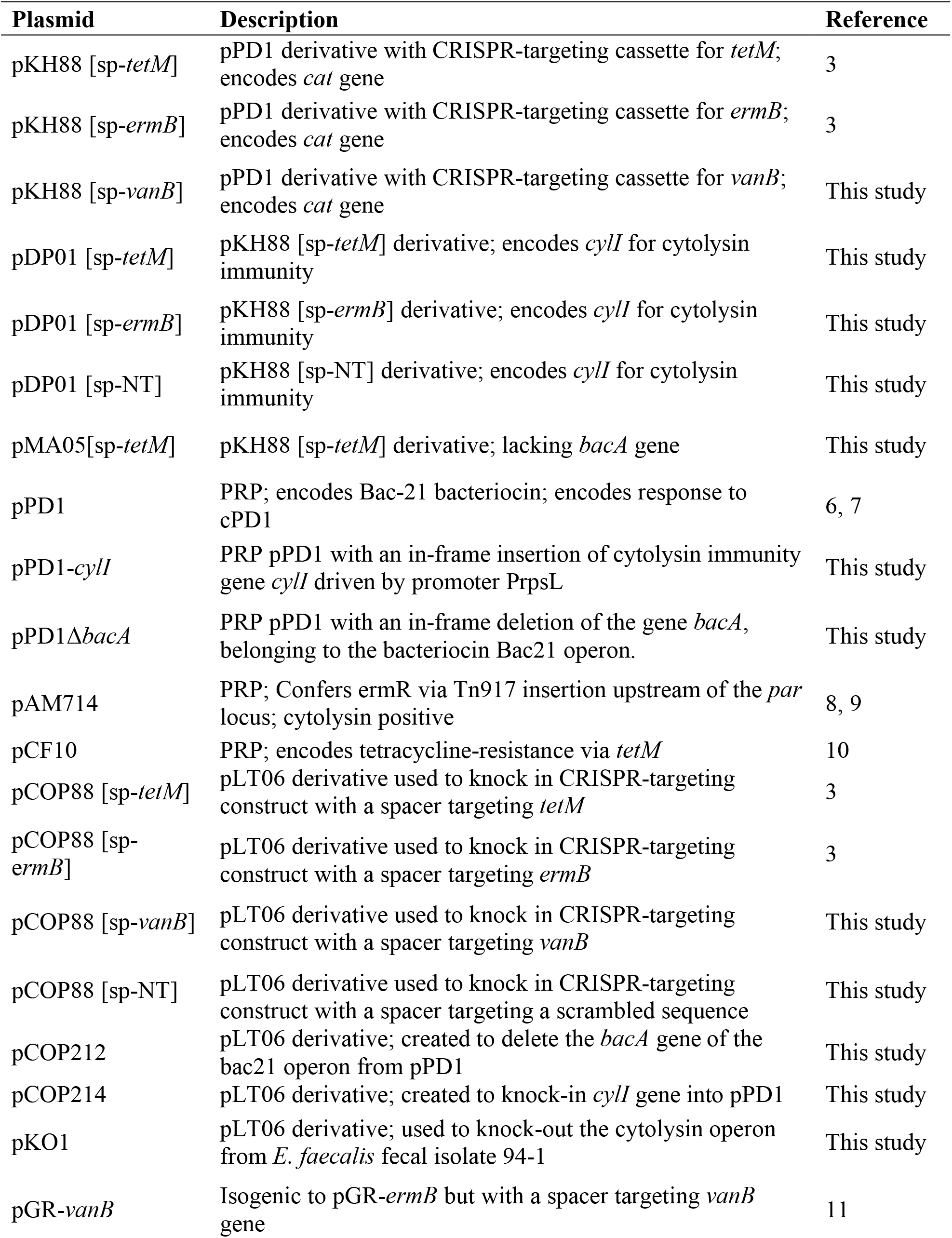
Plasmids used in this study.

**Table S4.**
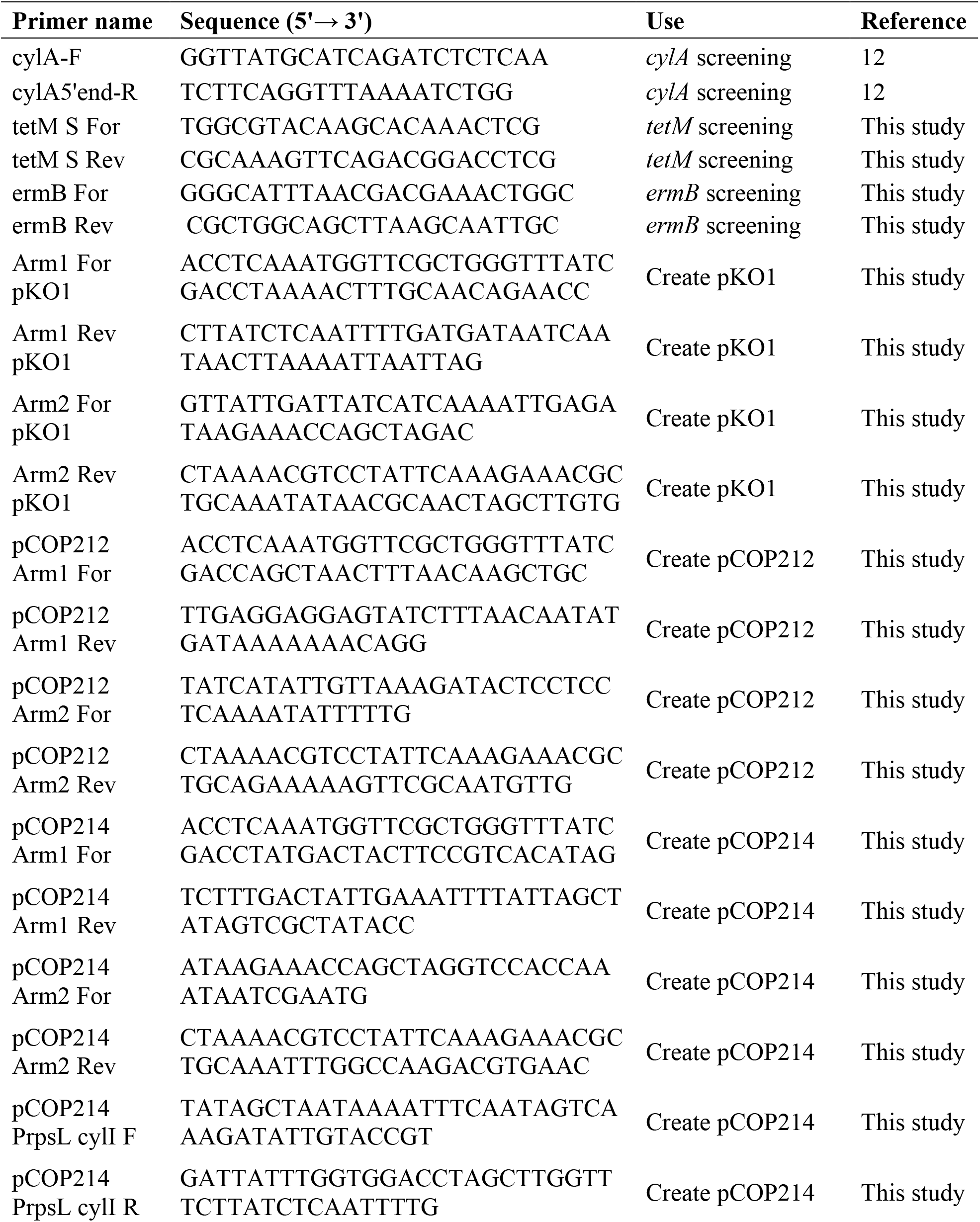
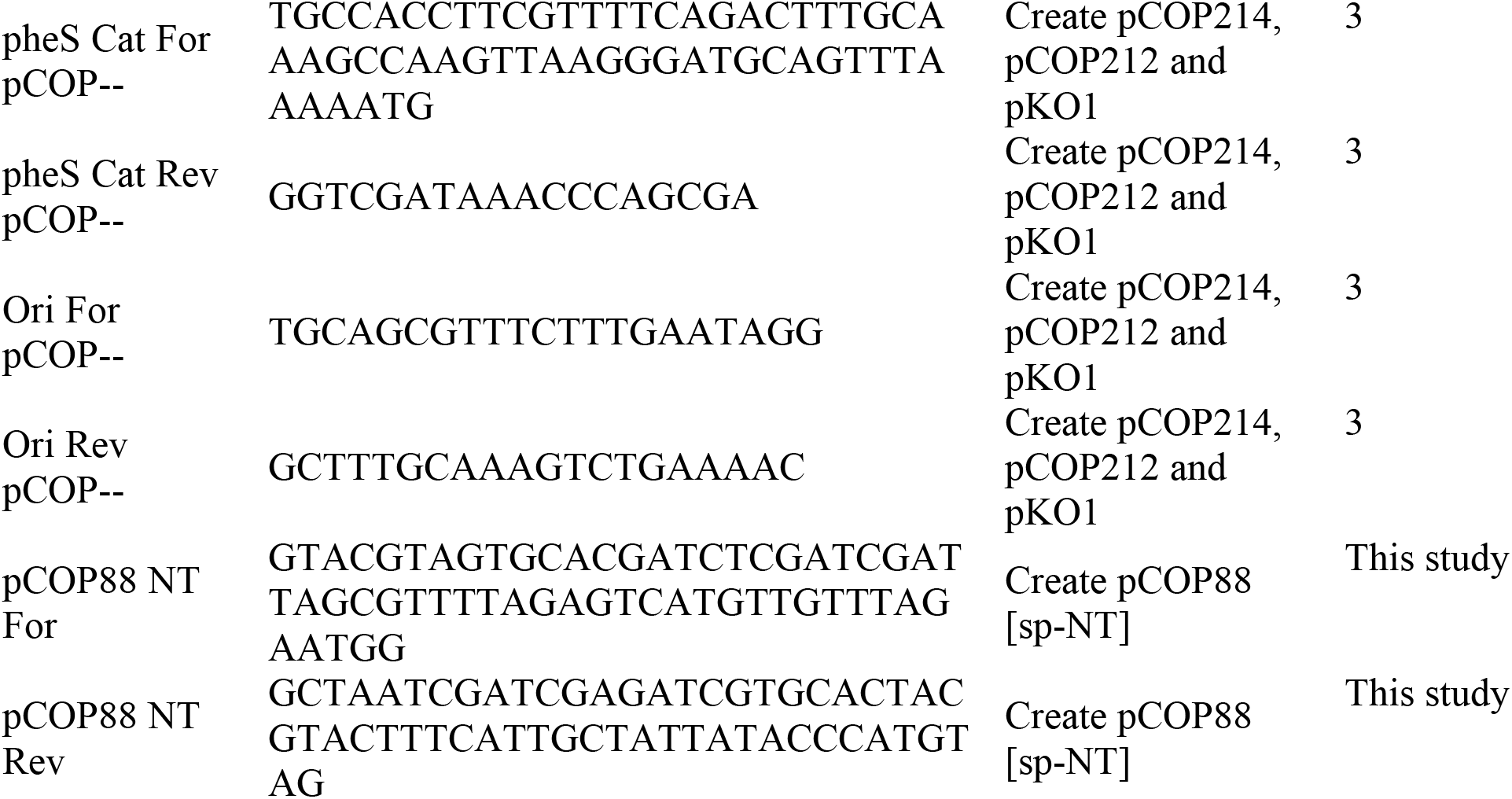
List of primers used in this study.

